# Host species and geography impact bee-associated RNA virus communities with evidence for isolation-by-distance in viral populations

**DOI:** 10.1101/2023.07.16.549238

**Authors:** Chris R. P. Robinson, Adam G. Dolezal, Irene L. G. Newton

## Abstract

Virus symbionts are important mediators of ecosystem function, yet we know little of their diversity and ecology in natural populations. The alarming decline of pollinating insects, especially the European honey bee, *Apis mellifera*, has been driven in part by worldwide transmission of virus pathogens. Previous work has examined the transmission of known honey bee virus pathogens to wild bee populations, but only a handful of studies have investigated the native viromes associated with these bees, limiting epidemiological predictors associated with viral pathogenesis. Further, social variation among different bee species might have important consequences in the acquisition and maintenance of bee-associated virome diversity.

We utilized comparative metatranscriptomics to develop a baseline description of the RNA viromes associated with wild bee pollinators and to document the interaction between hymenopteran insect social behavior and virus community composition. Our sampling includes five wild-caught, native bee species that vary in social behavior as well as managed honey bees. We describe 26 putatively new RNA virus species and contrary to our expectations, find few differences in virus diversity or abundance among socially variable bee species. Each bee species was associated with a specific virus community composition, even among sympatric populations of distinct host species. From 17 samples of a single host species, we recovered a single virus species despite over 600 km of distance between host populations and found strong evidence for isolation-by distance in associated viral populations. Our work adds to the small number of studies examining viral prevalence and community composition in wild bees.

## 2 Introduction

While long known to be important to human and agricultural health, virus symbionts are increasingly considered as critical modulators of ecosystem function [1, 2]. Despite their importance, virus ecology in natural populations of host species remains understudied. The co-evolving dynamics of virus-host interactions, shaped by myriad factors such as transmission rate, host density, or pathogen virulence, can be perturbed by minor changes in host ecology, resulting in rapid increases in virus transmission, virulence, and spread as evidenced by recent virus pandemics. Efforts to understand the impact of pathogen burden on ecosystems function, such as in the conservation of threatened species, necessitate a deeper understanding of virus ecology.

Among the most alarming population declines are those of pollinating insects, which provide essential ecosystem and agricultural services across many environments [3]. For most of the world, the European honey bee *Apis mellifera*, is the key managed pollinator, but wild bees have been shown to be important contributors to crop yield and agricultural function [4], as well as in their native ecosystems [3]. The worldwide decline in both native and non-native bee populations [5, 6] has been driven by a confluence of factors such as pathogen exposure, habitat loss, pesticide use [7] and pathogen exposure[7]. Among pathogens, the emergence of new or more widespread virus infections have garnered the most attention. In bees alone, there are over 50 described viruses [8, 9], many of which have been shown to have important implications for host survival [10]. However, despite the widespread and urgent concern about the role viruses play in the decline of bee populations worldwide, it remains that the majority of bee viral research has been conducted only in *A. mellifera*. Our understanding of virus ecology in other bee species, which often share overlapping pollinator networks with *A. mellifera*, is far more limited [11, 12]. A number of studies have examined the transmission of honey bee viruses to other bee species [6, 13, 14, 15], identifying evidence of virus pathogen transmission vectored by *A. mellifera*. While these studies have been important in describing the effects of virus infection and in illuminating pathogen networks among pollinating bees, they have focused almost exclusively on the viruses known to infect honey bees and have excluded investigating the native viromes of wild bee species. The absence of this data, addressed by only a handful of studies [9, 16] precludes the development of a baseline understanding of virus ecology in wild bee species and limits predictors of virus diseases in wild bees to the presence of sympatric honey bee populations.

Due to the many interactions intrinsic to pollinator networks, there are many potential opportunities for intra-and interspecific pathogen transmission, including the consumption of common resource stocks [17], vector-mediated transmission [18, 19, 20], and through sharing of the same floral resources [21, 13, 22]. Given the number of direct and indirect interactions between communities of pollinating bees, it is difficult to anticipate the degree of genetic and taxonomic diversity within bee-associated virus communities. Nearly all insect-infecting viruses are RNA viruses, which are generally characterized by relatively high mutation rates resulting from their lack of effective proofreading activity in their RNA polymerase [8]. This error-prone nature of RNA viruses can lead to high amounts of virus genetic diversity within a given host [8]. The fate of these genetic variants is then determined by ecological and evolutionary forces, such as selection, drift, and migration. Bee-associated RNA virus populations, undergoing weak to neutral selection, would be expected to be associated with high degrees of genetic diversity within the population. Low genetic diversity associated with RNA virus populations might be expected from positive selection on virus variants, such as from immune system responses to infection or arising from demographic changes, such as population bottlenecks originating from recent transmission events.

Inter- and intraspecific transmission of RNA virus population is expected to be a major source of virus infection among pollinating bees [21], and so we sought to investigate how the social behavior of bee species influences the genetic and taxonomic variation of associated virus communities. Species of pollinating bees can differ tremendously in behavior and the degree of sociality. Nearly all bees provide resources to the next generation of offspring, though how these resources are provided and distributed vary widely. Solitary bee offspring may receive only a single provisioning of resources to complete their development into adults [23] where as eusocial bees, such as *A. mellifera*, routinely share food between individuals and between developing offspring and might result in many more opportunities for the transmission of virus symbionts. Investigations of the bacterial communities of solitary bees as well as social bees reveal that the communities are often highly host-specific and consistent between individuals [24, 25]. Within other species of Hymenoptera, including the ants, the host-specificity of the associated bacterial community seems to be more relaxed [26].

Here, we conducted metatranscriptomic sequencing of five species of solitary and social halictid bees as well as the highly eusocial *Apis mellifera*. Our goal was to develop a baseline description of the RNA viromes associated with wild bee pollinators and to better understand the interaction between hymenopteran insect social behavior and virus communities. We chose to examine solitary and social halictid bee species because eusociality has evolved and been lost repeatedly within this group, resulting in a phylogeny of many closely related species demonstrating a spectrum of social behaviors [27, 24, 28, 29]. Samples from sympatric *A. mellifera* populations were included as a comparison since it represents a phylogentically distant eusocial species and is the most deeply studied bee species. We also investigated the impact of geography on associated RNA viromes by examining allopatric populations of a single solitary ground-nesting halictid, *Lasioglossum leucozonium*. Our motivating question asks whether the RNA viromes of social bees differ in composition or variation from the RNA viromes of closely related solitary bees. We hypothesized that RNA virome of social bees would experience reduced intraspecies variation due to the reduced environmental homogeneity and repeated interactions between social halictids, allowing for a higher number of horizontal transmissions. Finally, since most pollen foraging done by bee species is done within 3 km of their nest, we further hypothesized that allopatric populations of a single solitary halictid species, coupled with the intrinsically poor proofreading and mutation rates of RNA viruses, will lead to high divergence between populations.

## 3 Methods

### 3.1 Sample collection

Individual bee samples were collected from the Mid-Atlantic and Northeastern United States in 2016 and 2018 (Figure 1). A total of 38 samples were collected across six species of bee. The species *Augochlora pura*, *Apis mellifera*, *Agapostemon virescens*, *Lassioglossum versatum*, and *Augochlorella aurata* were collected from Princeton, New Jersey. *Lassioglossum leucozonium* was collected from several sites in the Northeastern US. These sites were Cobscook Bay, ME (44.839159, −67150384) Winter Harbor, ME (44.394613, −68.084561); Rangley, ME; (44.928539, −70.636231) Craneberry Lake, NY (44.204063, −74.831175); and Sunapee, NH (43.382356, −72.085406) (Supplemental Table #1).

**Figure 1:**
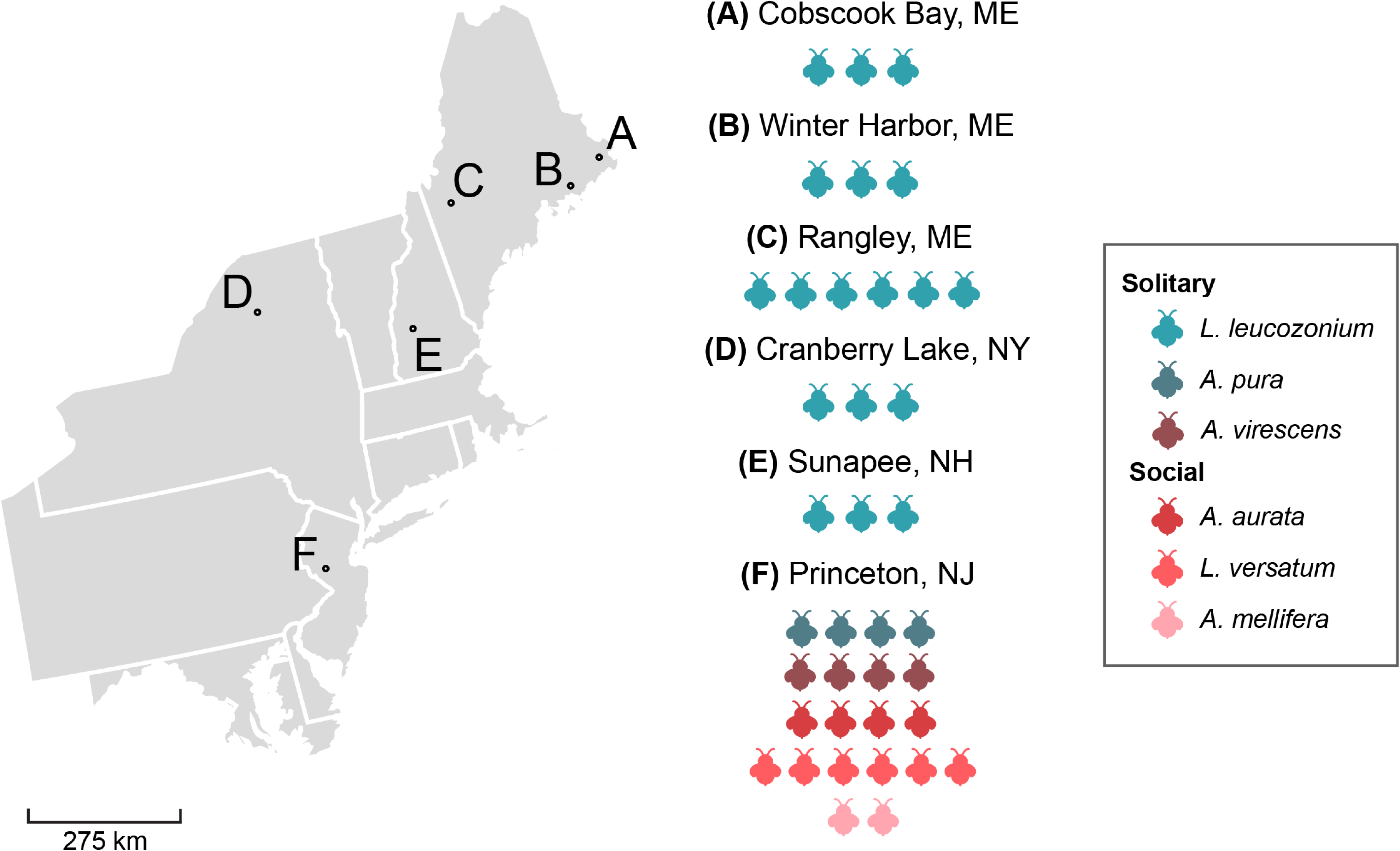
Locations of sample sites and species nomenclature. Letters on map correspond to individual sampling sites. The name of each sampling site and the number of bees collected at each site is to the right of the map. Color of bee represents bee species identity. The total number of bees icons associated with a given sampling site represents the number of bee samples collected at the site.

### 3.2 RNA library preparation and sequencing

RNA libraries were constructed from each individual sampled bee. The entire body of the bee was used for RNA extraction except for *A. mellifera* in which only the abdomen was used for extraction. In brief, individual bee samples were homogenized with sterile plastic pestles and treated with proteinase K. RNA was extracted using Zymo Quick-RNA Miniprep Plus kits. Host rRNA was removed using the NEBNext rRNA Depletion Kit (NEB #6310). Individual RNA libraries were created for each bee sample with NEBNext Ultra II Directional RNA Library Prep Kits. Each single-end 100 bp RNA library was sequenced on an Illumina NovaSeq 6000 at Princeton University across three flowcells.

### 3.3 RNA virus characterization, annotation, and discovery

Raw reads were quality checked with FastQC [30] and MultiQC [31] before trimming with Trimmomatic [32]. HiSat2 [33] was used to align reads from each individual RNA library to a corresponding reference genome. At the time of analysis, *L. versatum* did not have a reference genome and reads from these samples were aligned to the *L. leucozonium* reference genome. Except for *A. melifera*, reference genomes for all bee species were generated by the labs of Sarah Kocher and Erez Lieberman Aiden [34, 35, 36]. RNA reads that aligned to the reference were discarded, and the unaligned reads were used as input for assembly with rnavirusSPADES[37]. Assembled contigs were then used as inputs for CheckV [38] and Cenote-Taker2 [39] in order to assign initial confidence scores that a given contig was virus. We culled contigs that were either less than 500 base pairs or in which the presence of a virus hallmark gene could be confidently called. Viruses that passed this QC and maintained a sufficient virusity score were annotated with Cenote-Taker2.

Virus contigs were then screened for partial or complete RdRP sequences using annotations from Cenote-Taker 2 and Palmscan [40]. RdRP sequences were considered truly RdRP-associated if they were identified by both Cenote-Taker 2 and Palmscan or if Palmscan associated the sequence with a high confidence score. virus contigs containing a partial or complete RdRP sequence were then translated and clustered into family-equivalent OTUs using CD-HIT [41]. If two or more RdRP translations shared *>*40 % amino acid identity (AAI) over 80 % of the sequence, they were clustered into the same family-equivalent OTU. Contigs from each cluster were then then queried with BLASTP [42] in order to assign putative virus taxonomy. A virus species was considered novel if it shared *<*90 % RdRP amino acid identity (AAI) with known viruses in the non-redundant NCBI database [43].

### 3.4 Phylogenetic analysis of RNA viruses

Complete or partially complete RdRP sequences of all putatively novel and identified RNA viruses for phylogenetic analysis. The RdRP gene in RNA viruses is the standard in taxonomic comparisons between family-equivalent OTUs of RNA viruses, due to the conserved nature of its sequence relative to the sequence of other genes in the RNA virus genome [44]. Translated RdRP sequences were aligned using MAFFT [45] using default settings, and low quality aligned positions were trimmed with TrimAL with the *-automatic1* flag [46]. This alignment was then used to estimate maximum likelihood trees (ML) with IQTREE2 [47] with 1000 bootstrap replicates using the most appropriate model found by ModelFinder [48]. For family-level phylogenetic analysis, all known genomes associated with each virus family identified in this study were downloaded from NCBI RefSeq. Palmscan and custom bash scripts were used to identify and extract RdRP sequences in the NCBI virus genomes. RdRP sequences specific to each virus family, either from NCBI or from samples in this study, were aligned and trimmed with MAFFT and TrimAL before tree estimation with IQTREE2. A combination of bootstrap scores and SH-like branch support was used to validate tree topology of nodes in the estimated tree. In all trees the Astroviridae virus, Mamastrovirus 3, was used as outgroup.

### 3.5 RNA virus diversity and abundance

Absolute and relative virus abundances were estimated in each bee sample by mapping non-bee raw reads with bwa-mem [49] to each contig identified as virus. The number of mapped reads to a given contig were then normalized by the total number of reads in the sequencing library and the overall length of the contig as suggested by Roux et al [50].

The alpha diversity metrics, Shannon diversity, Simpson diversity, and richness were calculated among the samples at the species and family level using the Rhea script sets [51]. Comparing differences among samples separated by species or social behavior was done using a Tukey’s Honest Significant Difference Test with the R package agricolae [52]. We calculated beta diversity by quantifying virus diversity structure as composite variables along two axes of a principle coordinate ordination analysis and testing using Adonis tests from the vegan [53] package. These analyses were performed using R 4.2.2 and plotted with ggplot2 [54].

### 3.6 Fst calculations and analysis

To investigate the variation in RNA virus populations associated with allopatric populations of *L. leucozonium*, the cluster of contigs associated with *L. leucozonium* and from the family Narnaviridae were mapped with BWA-mem against the longest length contig within the cluster. Variant calling was done using LoFreq [55] without Indel calling. Variant calling files produced by LoFreq were used to calculate Nei’s *G_st_* (*F_st_*) (Equation 1) for each virus sub-population (e.g. single sample of bee).

We built a custom Python script to calculate *F_st_* (https://github.com/en-nui/BeeSocialityMetatranscriptomics). *F_st_* was calculated globally among all 18 Narnaviridae populations. Additionally, we calculated pairwise *F_st_* for each of the virus populations. Finally, each Narnaviridae population that shared a common sampling site (e.g. virus populations associated with a host that was sampled from the same sampling site) was grouped together as a single population, and we repeated the *F_st_*calculation for each of the geographically-defined virus populations. *F_st_* values in both analyses were then compared across a distance matrix, where pairwise Haversine distances (km) between sampling sites were compared to the corresponding differences in allele frequency variation using a Mantel test with the Pearson correlation method from the Vegan package.

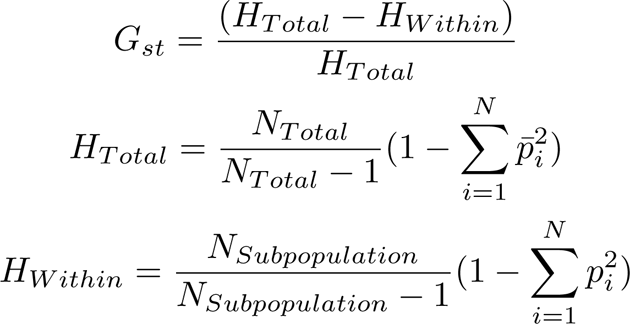

**Equation 1**. Calculations for Nei’s *G_st_* where the difference in average heterozygosity of the total population *H_T otal_* and a given subpopulation *H_W ithin_* is normalized by the average heterozygosity of the total population. *N_T otal_*and *N_Subpopulation_* are the total number of sampled chromosomes across all populations and within one subpopulation respectively. *P_i_* is the frequency of allele *i*.

## 4 Results

### 4.1 Characterization of known and novel viruses from bee hosts

We sequenced the RNA virus communities of 39 Apidae and Halictidae bees collected six locations around the Northeastern United States (Figure 1). Sequencing produced 1 574 309 491 reads and, after quality filtering, individual libraries specific to each species were mapped against corresponding host genomes and assembled into 156912 contigs. We chose a conservative approach to minimize false discovery and relied on stringent filtering of contigs by length and for known virus hallmark genes, reducing the total number of true virus contigs to 110. From these contigs, we recovered 9 putatively complete virus genomes. The majority of the remaining contigs contained mostly complete virus genomes and included a known RNA-dependent RNA polymerase or capsid gene.

From these 110 contigs, we characterized a total of 28 putative virus species OTUs from 12 different virus families. Clustering of virus OTUs was guided by recent comparisons of inter-and intrapopulation variation in virus populations [56] and by recommendations of the International Committee on Taxonomy of Viruses [57]. virus species were clustered by *>*95 % shared amino acid identity (AAI) over 80 % of the sequence length. virus families were clustered by *>*40 % AAI. BLASTP of all translated RdRP sequences resolved the species identity of 3 out of 28 species OTUs. The remaining 25 species OTUs, shared among all 6 species of sampled bee, were resolved only to the family level and are considered to be putatively novel virus species.

Families representing important honey bee virus pathogens, such as Dicistroviridae and Iflaviridae, were found in our survey with Iflaviridae appearing in both *Apis mellifera* and the solitary ground nesting bee, *Augochlora pura* (Figure 2a and 2b). Among halictid hosts, the abundance of viruses (detected by depth of read mapping to contigs) between host species, with *A. pura* and *Lasioglossum versatum* maintaining the highest levels of virus transcript compared to *Augochlora aurata*, and *Agapostemon virescens*. The virome of *A. virescens* was dominated equally by Reoviridae and Virgaviridae at levels much lower than halictids sampled in this study, and this may be a result of virus-infected plant tissue associated with pollen or plant debris consumed by the bee prior to RNA extraction.

**Figure 2:**
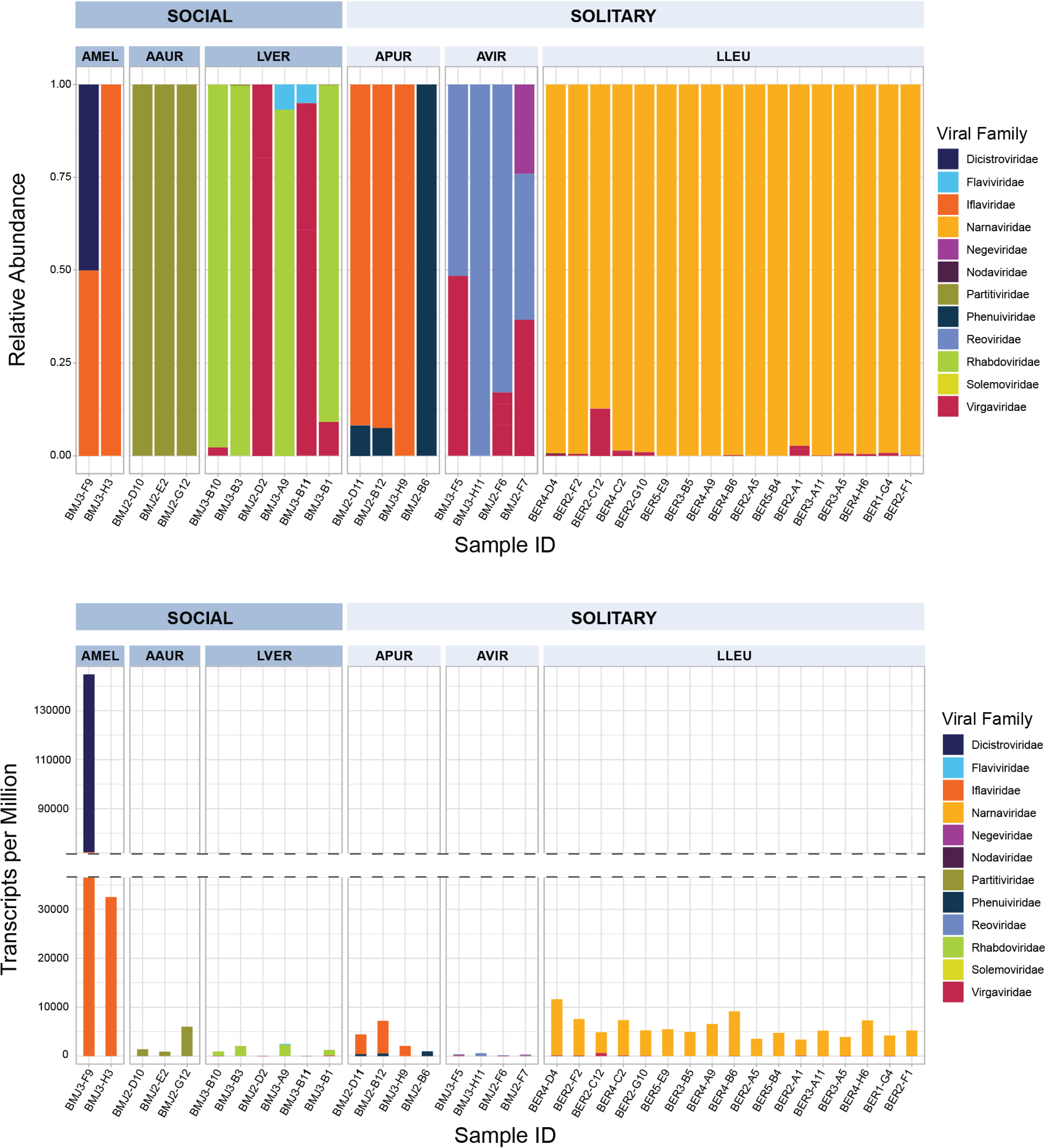
Relative abundance (2a; top) and absolute abundance (2b; bottom) of normalized RdRP virus transcripts across sampled bees in this study. Each bar plot along the X-axis represents an individual sampled bee and the Y-axis represents the proportion (top) or absolute number (bottom) of virus reads assigned to each family. Colors within each bar represent reads assigned to a given virus family.

Halictid virus transcripts were relatively low compared to extremely high levels of virus transcript in *A. mellifera* samples (Figure 2b, Fig S2). *A. mellifera* is known to persistently harbor viruses from Iflaviridae and Dicistroviridae [6]. Almost all bee-associated virus families characterized in this survey are known to infect insects, though we identified several virus species related to known plant pathogenic viruses In addition, a single species from the family Narnaviridae, uncommon outside of fungal hosts, dominates the virome of all sampled *Lassioglossum leucozonium*.

### 4.2 Novel RNA viruses

We identified novel species in 11 of the 12 virus families associated with sampled bees in this study (Figure 3). The majority of novel virus species fell into the family Virgaviridae which is primarily composed of plant pathogenic viruses (e.g. Tobacco Mosaic Virus). Virgaviridae species accounted for 9 of the 28 (32 %) of the novel RNA viruses identified in this study and were recovered from *A. virescens*, *L. versatum*, and *L. leucozonium*. Individual *L. leucozonium* samples harbored high Virgaviridae diversity and were associated with 6 of the 9 novel Virgaviridae species. The first of these species (Figure 4, Group 1) is most closely associated with Gentian Ovary Ringspot virus (YP 009047252.1), a virus first isolated in 2014 and is vectored by pollinating insects [58]. Long branch lengths are indicative of high levels of divergence, though clustering by amino acid identity of known and novel virus RdRPs place this clade into the genus Goravirus. The second Virgaviridae species (Figure 4, Group 2) identified in *L. leucozonium* shares the highest identity with Lychnis ringspot virus (YP 009508258.1), a plant pathogen of dicot plants. Amino acid clustering of all RdRPs in this clade fall into the genus Hordeivirus, which are known to be transmitted by both pollen and seed [59]. The remaining 6 novel species (Figure 4, Groups 3 - 5) share high sequence divergence with one another and with the closest known relative, Nephilia clavipes virus 3 (YP 009552459.1), recovered from the Golden Orb-Weaver spider [60]. Virgaviridae transcripts were uniformly low in all isolates, and it is difficult to distinguish whether the association of these virus species is due to infection of bee body issue, or if they were associated with plant debris on or consumed by bees during foraging. In the case of this study, the 9 novel Virgaviridae species are most likely associated with the pollen foraged or consumed directly by these three bee species and reflect the significance of pollinators as vectors of plant pathogenic viruses.

**Figure 3:**
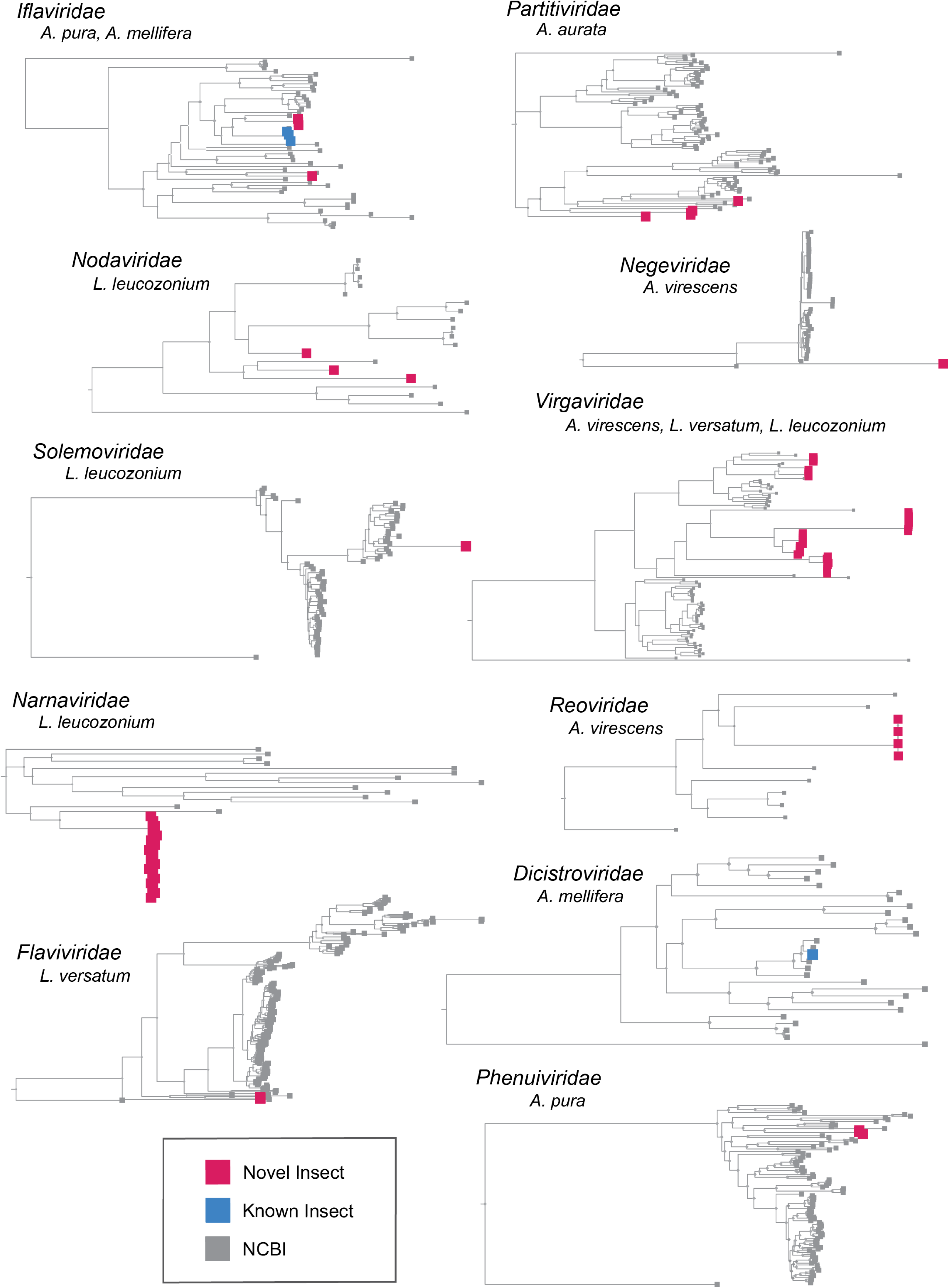
Overview of viruses found in bees sampled in this study. Bee hosts associated with a virus family are listed underneath the corresponding family name (Rhabdoviridae excluded). Viruses associated with this study are separated into three groups. Novel insect-associated viruses are depicted by red boxes. Known insect-associated viruses are depicted by blue boxes. Finally, virus genomes taken from NCBI RefSeq are depicted by gray boxes.

**Figure 4:**
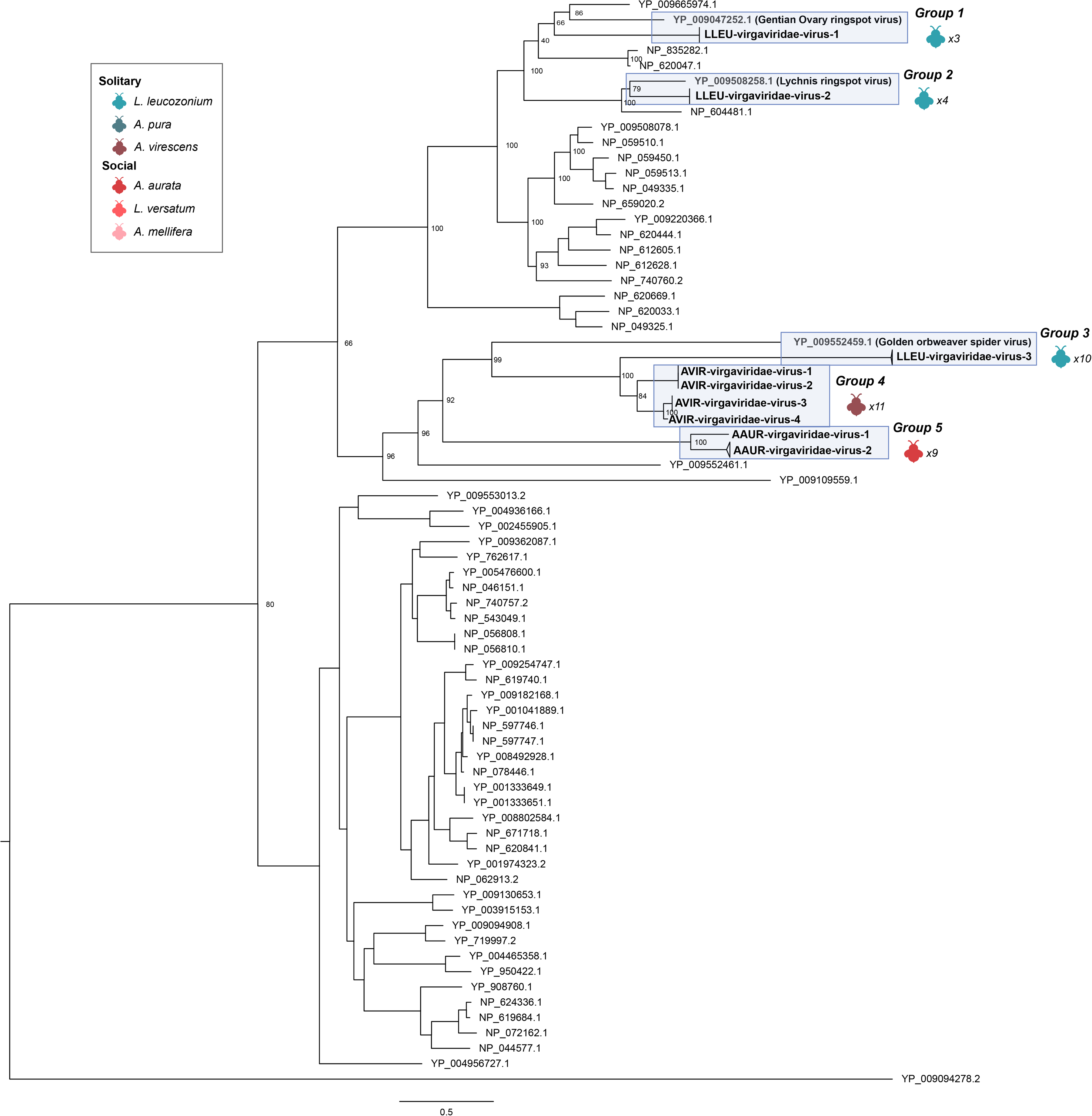
RdRP phylogeny of Virgaviridae viruses. Tree is midpoint rooted for clarity only. Viruses sampled in this study are highlighted in blue. Bootstrap values from 1000 replicates for select nodes are provided. Bee icon color represent species host that virus group was associated with; number represent the number of bee samples associated with virus group. The scale bar represents the number of amino acid substitutions.

Each of the 5 halictid species harbored novel virus species that were unique to each halictid species host. The viromes of the social halictid *A. aurata* were dominated by a single species of Partitiviridae, (Fig S3, clade 1). RdRP sequences within this species were most closely related to Cryptosporidium pavum 1 virus (YP 009508065.1) [61], though external long branches of this clade suggest significant sequence divergence. Likewise, the other social halictid sampled in this survey, *L. versatum*, was associated with Virgaviridae and a single Rhabdoviridae and Flaviviridae species. The number of assigned Rhabdoviridae transcripts in these samples were much higher than the number of transcripts assigned to Virgaviridae, suggesting that Rhabdoviridae association in *L. versatum* reflects an active virus infection (Fig S4). The solitary halictids harbored a higher absolute number of virus species compared to the social halictids, though this difference in diversity was not statistically significant. We identified 2 novel species of Iflaviridae (discussed further in the text) and 2 species of Phenuiviridae across all 5 samples. This species of Phenuiviridae was most similar to the Mothra mobuvirus (YP 009666266.1) (Fig S5), and RdRP sequence clustering places both viruses within the genus Mobuvirus. The final solitary halictid sampled from Princeton, *A. virescens*, harbored 4 novel species of virus across Negeviridiae (Fig S6), Reoviridae (Fig S7), and Virgaviridae. RdRP sequences in these samples were highly divergent and showed no close association with any known ssRNA viruses in NCBI RefSeq (Figure 3). virus transcripts in these samples were incredibly low and were similar to other Virgaviridae transcript levels in other sampled bee isolates, suggesting that these viruses are passively associated with environmental debris associated with or consumed by *A. virescens* during sampling.

The solitary halictid, *L. leucozonium*, sampled across 5 allopatric populations, showed a surprising trend. As discussed above, individual samples of *L. leucozonium* were associated with a high diversity of Virgaviridae species, possibly reflecting a diversity of pollen samples consumed by this species during foraging. In contrast to this diversity of plant pathogenic viruses, all 18 samples of *L. leucozonium* were infected by a single species of Narnaviridae. This species is most closely related to Saccharomyces 20S RNA narnavirus (NP 660178.1) (Figure 5).

**Figure 5:**
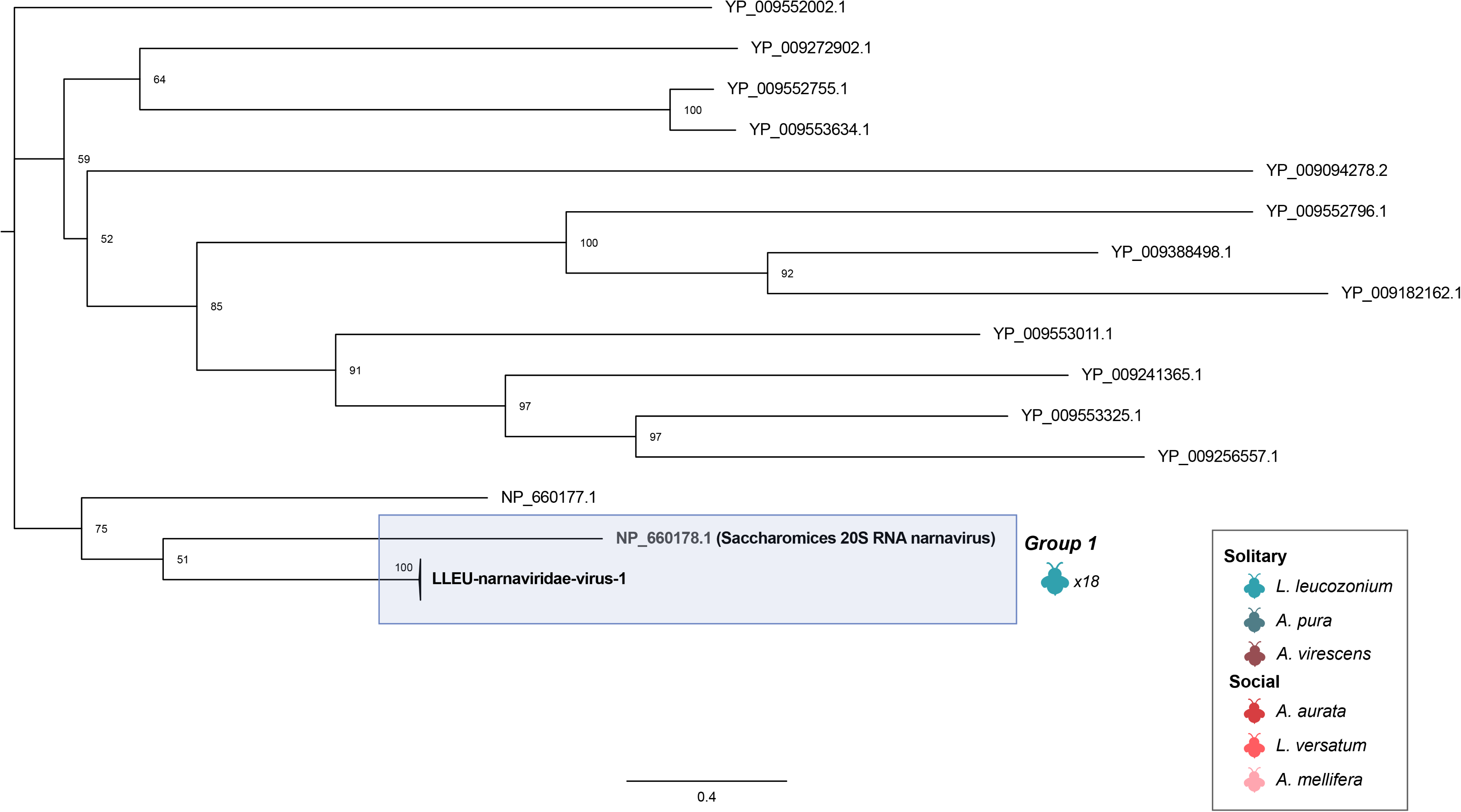
RdRP phylogeny of Narnaviridae viruses. Tree is midpoint rooted for clarity only. Viruses sampled in this study are detailed highlighted in blue. Bootstrap values from 1000 replicates for select nodes are provided. Bee icon color represent species host that virus group was associated with; number represent the number of bee samples associated with virus group. The scale bar represents the number of amino acid substitutions.

### 4.3 No evidence of Iflaviridae transmission by domesticated bees to wild bees

Species-level OTU clustering of RdRP protein sequences revealed 3 virus species distributed across *A. mellifera* and *A. Pura* (Figure 6). Though Iflaviridae have been shown to widespread virus symbionts of insects [62], previous work has shown that viruses in this family, particularly Deformed Wing Virus, are a major cause of the worldwide decline in honey bee populations [63, 64, 65] and are important pathogens of other pollinators such as bumblebees [66]. BLASTP of the RdRP protein sequence identified that Deformed Wing Virus, one of the 3 species of Iflaviridae (Figure 6, Group 2) found in our survey, was found in both of the honey bee samples, and combined with the extremely high level of virus transcript found in these samples (Figure 2b), is indicative of active virus infection – possibly a result of feeding by the parasitic mite, *Varroa destructor* [65].

**Figure 6:**
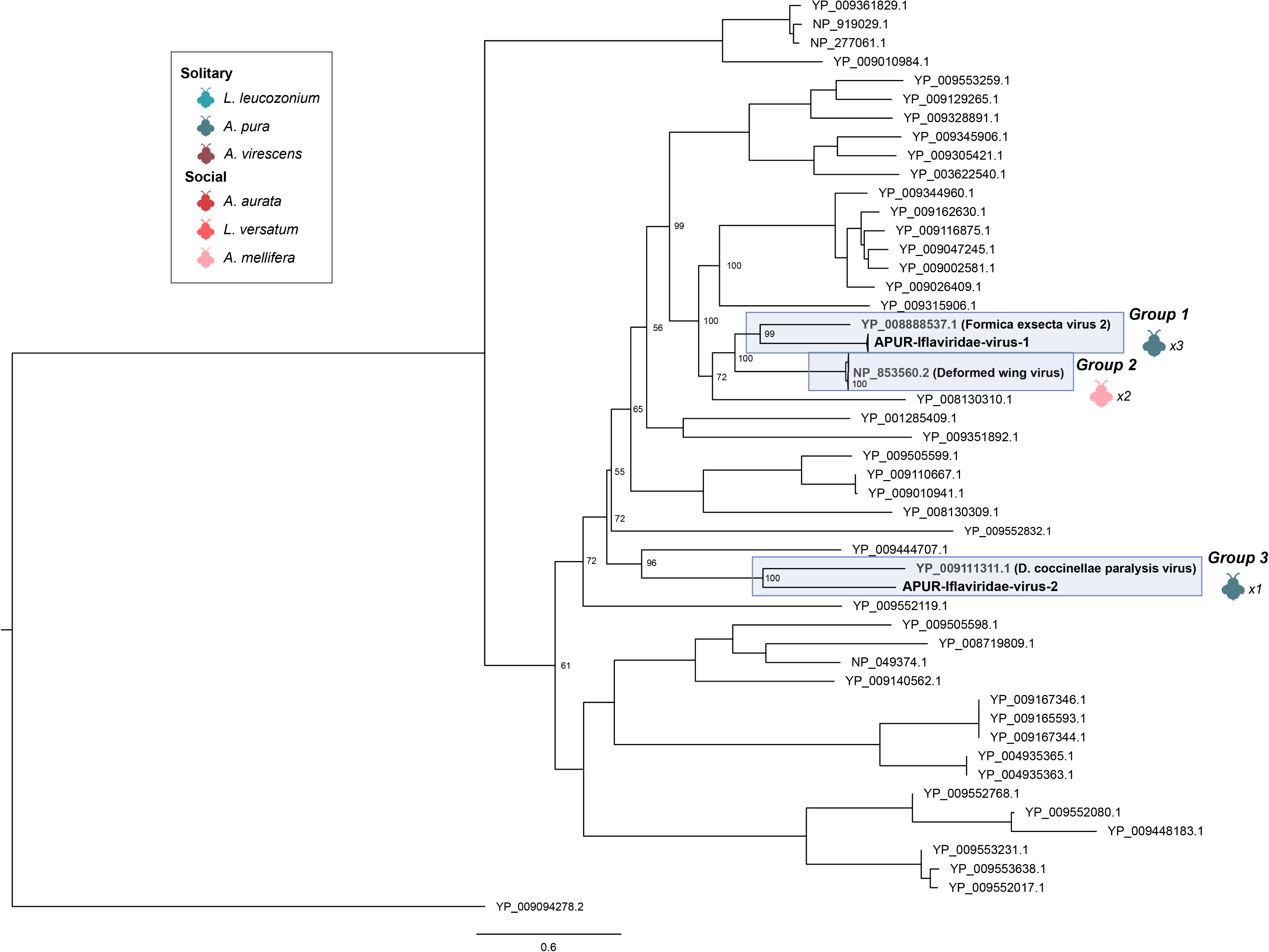
RdRP phylogeny of Iflaviridae viruses. Tree is midpoint rooted for clarity only. Viruses sampled in this study are highlighted in blue. Bootstrap values from 1000 replicates for select nodes are provided. Bee icon color represent species host that virus group was associated with; number represent the number of bee samples associated with virus group. The scale bar represents the number of amino acid substitutions.

Further, we identified two novel virus species in Iflaviridae. Phylogenetic analysis of shared virus RdRPs in Iflaviridae revealed that a virus species infecting APUR, (APUR-Iflaviridae-1 through APUR-Iflaviridae-3), is most closely related to Formica exsecta virus 2 (Figure 5, Group 1), a virus associated with the ant *Formica exsecta* [67]. Long branch lengths separating the Formica exsecta 2 virus and APUR-IFLAVIRIDAE-1 suggest relatively long periods of divergence. Members of the clade occupied by Deformed Wing Virus formed the next most related group. The final novel Iflaviridae virus species, APUR-IFLAVIRIDAE-2, was found to be most similar to the virus Dinocampus coccinellae paralysis virus (Figure 5, Group 3). This virus has been shown to induce behavior manipulation, such as tremors and paralysis, of ladybird beetles that have been parasitized on by the wasp, *Dinocampus coccinellae* [68]. Similar to above, long branch lengths between this virus and APUR-IFLAVIRIDAE-2 indicate high levels of divergence.

### 4.4 Evidence for isolation-by-distance in wild bee-associated viral populations

*L. leucozonium* hosts sampled from all 5 sample sites were actively infected by viruses from the family Narnaviridae. Despite over 600 kilometers of distance between some sites and the low replication fidelty intrinsic to RNA viruses, RNA virus populations in these samples are remarkably similar (phylogeny figure). Each sample of *L. leucozonium* is dominated by a putatively novel virus in the virus family *Narnaviridae*. Membership in this family was assigned by comparing the translated nucleotide identity (AAI) of the virus gene encoding RNA-dependent RNA polymerase (RdRP). RdRP sequences from all *L. leucozonium*-associated Narnaviridae shared 95 % amino acid identity over 80 percent of the sequence length, indicating that Narnaviridae found in each *L. leucozonium* belongs to the same species (Fig 5). Viruses within Narnaviridae have positive-sense single-stranded RNA genomes of 2.3 to 2.9 kb [69] and encode a singular RdRP protein. We recovered a complete Narnaviridae genome from each individual *L. leucozonium* with an average length of approximately 3.2 kb.

The presence of the same virus species among all sampled *L. leucozonium* individuals allowed us to compare the variation in allele frequency of sub-populations of Narnaviridae (e.g., one sampled bee host) to the variation in allele frequency of the entire population of RNA viruses (e.g. between all sampled bee hosts across all sample sites). To explore this variation, we developed a custom Python script to calculate Nei’s Gst [70] for pairwise *F_st_* comparisons between each single Narnaviridae population and Narnaviridae populations from each sampling site. We calculated a global *F_st_* of 0.484, which is indicative of high interpopulation genetic variation. Results from the individual pairwise *F_st_* calculations produced *F_st_* values indicating high levels of population differentiation with an average *F_st_* of 0.322 (Fig S9). Though there was significant amount of differentiation between individual Narnaviridae populations, a Mantel test using Spearman’s rank correlation found no association with pairwise *F_st_* and Haversine distance (*p* = 0.607).

Pairwise *F_st_* values produced by comparing Narnaviridae associated with geographically-defined hosts showed a contrasting trend. Calculations of pairwise *F_st_*between geographically-defined Narnaviridae populations produced an average *F_st_* of 0.046 and a maximum *F_st_* of 0.089, indicative of far less differentiation between geographically-distinct populations of Narnaviridae than between individual populations (Figure 7). Further, we found that geographic distance contributes significantly to observed genetic differentiation between these populations. A Mantel test using Spearman’s rank correlation, including Haversine distance and values of pairwise *F_st_*, was indicative of a significant correlation between geographic distance and genetic differentiation, (*r* = 0.5988 P = 0.025). Further, generating a scatterplot of geographic distances between any two populations showed a continuous cline of *F_st_*. These results suggest that as distance increases between any two geographically-defined populations of hosts, there is an increase in the genetic differentiation between populations of associated Narnaviridae (Figure 7).

**Figure 7:**
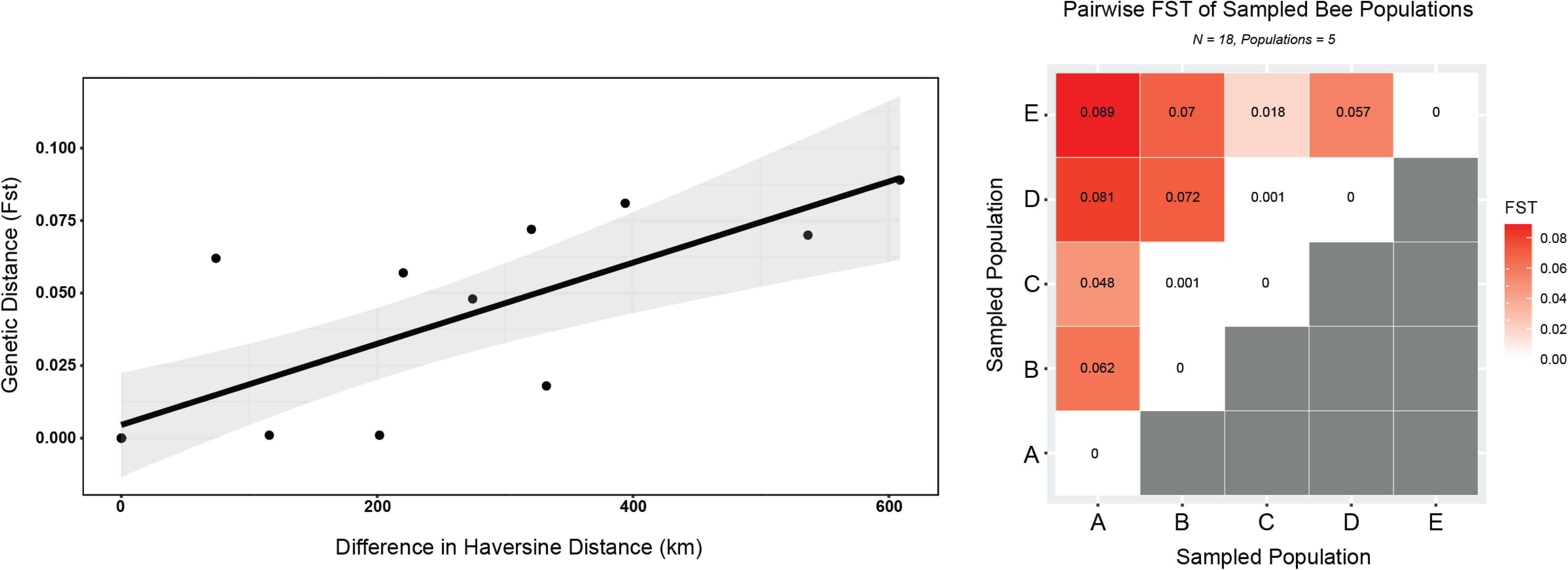
Scatterplot depicting pattern between pairwise Haversine distance (km) and *F_st_* (7a; left). Individual points represent populations of Narnaviridae associated with bee hosts sampled from the same site. A Mantel test produced a significant correlation between the Haversine distance between any two sampling sites and pairwise *F_st_* (*p* = 0.025). Plotted linear regression is not representative of the Mantel test but is for clarity only. A matrix of pairwise *F_st_* comparisons representing patterns of genetic differentiation among Narnaviridae populations associated with 5 geographically-defined sites (7b; right). Letters on the X-Axis correspond to sampled *L. leucozonium* population (Figure 1). Color represents the degree of genetic differentiation between any pair of combinations. Darker hues indicates higher values of differentiation. Genetic differentiation is defined by a scale on the right side of the figure.

### 4.5 Host species and not social behavior explains virome composition and diversity

Except in the case of *L. leucozonium*, no two halictid species of bee hosts shared the same virus species or family. Indeed, virus families were species-specific except for Virgaviridae, and so differences in comparing the virus communities associated with social and solitary samples might reflect peculiarities in aspects of virus biology. With this context in mind, we analyzed the intra-sample virus diversity and richness. We first tested for differences in the number and diversity of virus OTUs (species and families) associated with behaviors of halictid bees. When comparing all halictid bees sampled from the same locality (Princeton), we found no difference in the virus Richness or Shannon and Simpson indices (Richness: p = 0.585, Shannon: p = 0.184, Simpson: p = 0.156) (Figure S10). We also compared differences in diversity of virus communities by including *A. mellifera* (also sampled from the same locality) and *L. leucozonium* (which was geographically dissimilar), but found no differences in effect (Richness: p = 0.5, Shannon: p = 0.201, Simpson: p = 0.195). We then investigated the effect of species association on virome richness and diversity. As above, we analyzed only the halictid hosts sampled from the same locality. Analysis via ANOVA and post-hoc analysis with Tukey’s Honest Significance Test (HSD) found no significant difference in Richness or Shannon and Simpson indices (Richness: p = 0.265, Shannon: p = 0.243, Simpson: p = 0.207) (Figure S11).

Despite no significant difference in virus species diversity and richness between different sympatric species of bee host, our sampling showed that distinct virus families were present in each of the different bee species and that species varied significantly in their virus species composition (Figure 2a, Figure 2b). Principal coordinate analysis using Bray-Curtis dissimilarity of the relative abundances of associated virus OTUs distributed the viromes into distinct clusters at both the species and family level (Adonis results: DF = 5, *R*^2^ = 0.824, P = 0.00001) (Figure 8). Bees sampled from the same locality clustered closer together in ordinate space than with bees sampled from other localities, such as *L. leucozonium*), though virus OTUs from *L. versatum* (also sampled from the same locality) showed high dissimilarity. These findings are consistent with other studies investigating interactions between sociality and microbiome composition in bees [24, 25, 71, 72, 73, 74].

**Figure 8:**
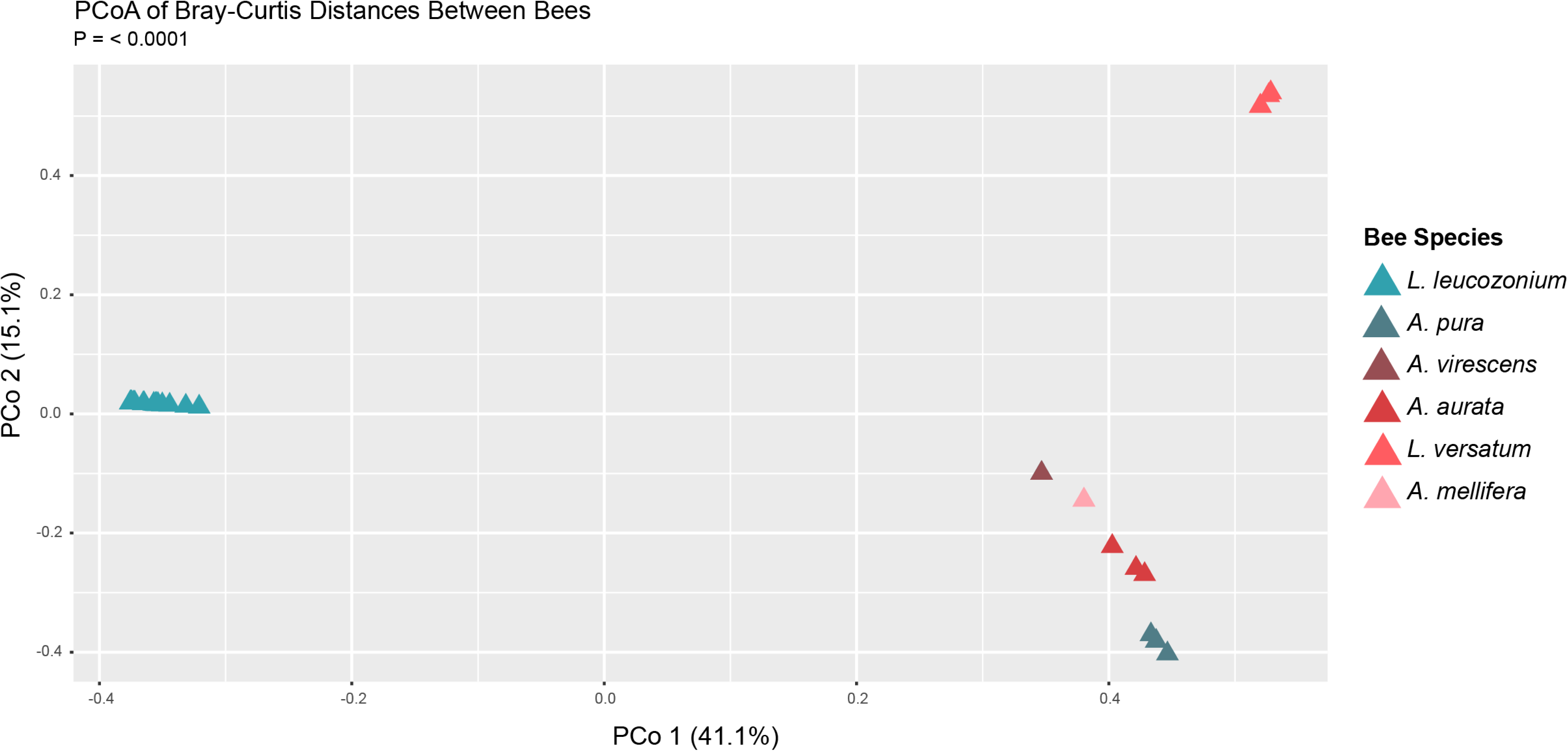
Bray-curtis dissimilarity Principle Coordinate Analysis based on absolute abundance of normalized transcript for each virus species OTU described in this study. Each triangle represents an individual bee sample. Color depicts the associated bee species. PERMANOVA revealed that overall virus species composition was significantly distinguished by host species. (P = 0.00001)

## 5 Discussion

Here, we have used comparative metatranscriptomics to investigate the viromes of 6 species of bee, including 5 native halictid bees, across a spectrum of host sociality. Our sampling includes five wild bee species, drawn from both sympatric and allopatric populations, and our analysis identified 26 putatively novel virus species across 12 virus families. The majority of the novel virus species fell into the family Virgaviridae, underlining the importance of pollinators as vectors of pollen-born plant pathogenic viruses. virus species from the family Iflaviridae, containing some of the most virulent and pathogenic viruses affecting honey bees, appeared in samples of both honey bee and the wild *A. pura*. Though we found no evidence of recent transmission of honey bee-related viruses in *A. pura*, our study is one of only a handful of studies that has surveyed the viromes of wild bees and examined signatures of virus spillover. The ongoing virus pandemic affecting the honey bee, exacerbated by the parasitic mite, *Varroa destructor*, has important ecological implications for wild bees, as infected apiaries may serve as reservoirs for virus transmission. Overall, virus transcripts associated with bees in our study were relatively low and may reflect a healthy equilibrium between virus and host. In contrast, sampled honey bees were associated with high transcript levels of Deformed Wing Virus and Israeli Acute Paralysis Virus. These results corroborate earlier investigations of RNA viral transcript levels across many bee taxa [6].c

Though much is known about virus ecology in social bees (A. mellifera), we sought to investigate the importance of sociality in acquisition and maintenance of virus communities in wild bees. Comparing populations of social bees to solitary bees, recent studies [24, 25, 72] have found little evidence for the influence of sociality on associated bacterial communities in wild, non-corbiculate bees. Though solitary bees in our study were associated with a larger number of virus species, comparisons to closely-related and sympatric social species revealed no effect of sociality on virus diversity or abundance. Shell and Rehan [25] did find significant differences when examining the effect of sociality on intrapopulation metagenomic variation, and it is likely that our approach to RNA virus taxonomy (e.g. assigning species at 95 % AAI) is too broad. Similar to the studies above, we found evidence for strong species-specific association with RNA viruses (Figure 8). Despite similar ecology and nesting strategies, sympatric populations of 4 halictid species were associated with distinct RNA virus families. Further, five allopatric populations of a single species, *L. leucozonium*, were all associated with the same species of Narnaviridae. It is unclear how *L. leucozonium* is associated with Narnaviridae, and it is surprising how little divergence has occurred despite the low replication fidelity of RNA viruses and over 600 kilometers of distance between *L. leucozonium* population in this study. Further work may investigate the presence of an endogenous retrovirus associated with *L. leucozonium* and the potential transcription of the retrovirus-associated loci.

The presence of Narnaviridae in all isolates of *L. leucozonium* allowed us to investigate the amount of virus diversity associated with allopatric populations of *L. leucozonium*. Two studies [25, 72] found that the microbiome of select wild bees species is influenced by its local environment. We investigated this and found significant evidence for Isolation-By-Distance in Narnaviridae metapopulations. Though individual pairwise comparisons of Narnaviridae population diversity found no association with Haversine distance, these findings suggest that Narnaviridae populations associated with individual *L. leucozonium* hosts found within the same geographic locality are as different from one another as they are from sampled hosts more than 600 kilometers away. This high population differentiation can be explained by the low fidelity rates shared by most RNA viruses [8] and by little to no migration between individual virus populations. Additionally, these virus populations may be a result of recent infection, and the high amount of genetic diversity within these populations may reflect a history of recent population expansion. Treating all Narnaviridae populations within a single sampling site as a metapopulation revealed a significant correlation between genetic and Haversine distance. This association may reflect signatures of host-virus coevolution as higher relatedness between sympatric populations of *L. leucozonium* might increase the frequency of alleles of specific immune genes and lead to less varied virus populations. Further work comparing loci associated with known immune genes in geographically-distinct hosts might provide support for this hypothesis.

Given the importance of RNA viruses to pollinator health, it is critical that we understand the interactions between wild bee species and their virus symbionts. Though virus research in the honey bee system is undoubtedly important and has taught us much about Hymenoptera-virus interactions, it is becoming increasingly clear that interactions and potential mechanisms between microorganisms and their bee hosts are species-specific. However, it is surprising the consistency of this specificity regardless of host behaviors or geography. Eusocial organisms have long been a fascination and special difficulty of evolutionary biology [75], and here we provide no evidence that social behavior correlates with any differences in diversity or composition between the viromes of social and solitary bees. However, the relatively simple viromes of this socially plastic family of bees, in addition to our understanding of the honey bee metagenome, incites further study in understanding the relationship between host sociality and symbiosis.

## Supporting information

Suupelemental Data

## 6 Acknowledgements

We thank Dr. Benjamin E. R. Rubin for sample collection, library creation, sequencing, and for sharing the data – all of which was supported by a USDA NIFA Postdoctoral Fellowship (2018–67012-28085). We thank Dr. Sarah Kocher and her lab for helping with sample collection, sequencing, and sharing the unpublished genomes. Genome assemblies and sequencing data for *L. leucozonium*, *L. versatum*, *A. aurata*, *A. pura*, and *A. virescens* are used with permission from the DNA Zoo Consortium (dnazoo.org) and from Dr. Sarah Kocher [34]. We are grateful to Aya McGee for her contributions to figure design and annotation, and we are grateful to Dr. Ĺılian Caesar for her suggestions and edits. This work was financially supported by a Costco/Project Apis m. research grant (to CR) and an NSF IOS Collaborative Research grant (2005306) and NSF DBI Biology Integration Institutes grant (2022049) to ILGN.

## 7 Data Availability

Metagenomic datasets for each bee sample have been deposited into the NCBI SRA database. The accession numbers for these samples are SAMN33439837 - SAMN33439874 under the BioProject PR-JNA938525. Scripts for the *F_st_* calculations are available on GitHub.

## References

[1] Chelsea L Wood and Pieter TJ Johnson. “A world without parasites: exploring the hidden ecology of infection”. In: Frontiers in Ecology and the Environment 13.8 (2015), pp. 425–434.

[2] K Eri Wommack et al. “Counts and sequences, observations that continue to change our understanding of viruses in nature”. In: The Journal of Microbiology 53.3 (2015), p. 181.

[3] Alexandra-Maria Klein, et al. “Importance of pollinators in changing landscapes for world crops”. In: Proceedings of the royal society B: biological sciences 274.1608 (2007), pp. 303–313.

[4] Lucas A Garibaldi, et al. “Wild pollinators enhance fruit set of crops regardless of honey bee abundance”. In: science 339.6127 (2013), pp. 1608–1611.

[5] Laura A Burkle, John C Marlin, and Tiffany M Knight. “Plant-pollinator interactions over 120 years: loss of species, co-occurrence, and function”. In: Science 339.6127 (2013), pp. 1611–1615.

[6] Adam G Dolezal, et al. “Honey bee viruses in wild bees: viral prevalence, loads, and experimental inoculation”. In: PloS one 11.11 (2016), e0166190.

[7] Simon G Potts, et al. “Global pollinator declines: trends, impacts and drivers”. In: Trends in ecology & evolution 25.6 (2010), pp. 345–353.

[8] Raul Andino and Esteban Domingo. “Viral quasispecies”. In: Virology 479 (2015), pp. 46–51.

[9] David J Pascall, et al. “Virus prevalence and genetic diversity across a wild bumblebee community”. In: Frontiers in Microbiology 12 (2021), p. 650747.

[10] Dino P McMahon, et al. “Emerging viruses in bees: From molecules to ecology”. In: Advances in virus research 101 (2018), pp. 251–291.

[11] Brittany Elliott, et al. “Pollen diets and niche overlap of honey bees and native bees in protected areas”. In: Basic and Applied Ecology 50 (2021), pp. 169–180.

[12] Ingolf Steffan-Dewenter and Teja Tscharntke. “Resource overlap and possible competition between honey bees and wild bees in central Europe”. In: Oecologia (2000), pp. 288–296.

[13] Peter Graystock, Dave Goulson, and William OH Hughes. “Parasites in bloom: flowers aid dispersal and transmission of pollinator parasites within and between bee species”. In: Proceedings of the Royal Society B: Biological Sciences 282.1813 (2015), p. 20151371.

[14] Dino P McMahon et al. “A sting in the spit: widespread cross-infection of multiple RNA viruses across wild and managed bees”. In: Journal of Animal Ecology 84.3 (2015), pp. 615–624.

[15] Peter Graystock, Dave Goulson, and William OH Hughes. “The relationship between managed bees and the prevalence of parasites in bumblebees”. In: PeerJ 2 (2014), e522.

[16] Karel Schoonvaere, et al. “Study of the metatranscriptome of eight social and solitary wild bee species reveals novel viruses and bee parasites”. In: Frontiers in microbiology 9 (2018), p. 177.

[17] Yan Ping Chen, et al. “Israeli acute paralysis virus: epidemiology, pathogenesis and implications for honey bee health”. In: PLoS pathogens 10.7 (2014), e1004261.

[18] CE Burnside. “The cause of paralysis of honeybees”. In: Am. Bee J 85 (1945), pp. 354–363.

[19] BV Ball. “Varroa jacobsoni as a virus vector”. In: Present status of varroatosis in Europe and progress in the Varroa mite control (1989), pp. 241–244.

[20] PL Bowen-Walker, SJ Martin, and A Gunn. “The Transmission of Deformed Wing Virus between Honeybees (Apis melliferaL.) by the Ectoparasitic MiteVarroa jacobsoniOud”. In: Journal of invertebrate pathology 73.1 (1999), pp. 101–106.

[21] Rajwinder Singh, et al. “RNA viruses in hymenopteran pollinators: evidence of inter-taxa virus transmission via pollen and potential impact on non-Apis hymenopteran species”. In: PloS one 5.12 (2010), e14357.

[22] Samantha A Alger, P Alexander Burnham, and Alison K Brody. “Flowers as viral hot spots: Honey bees (Apis mellifera) unevenly deposit viruses across plant species”. In: PLoS One 14.9 (2019), e0221800.

[23] Cécile M Antoine and Jessica RK Forrest. “Nesting habitat of ground-nesting bees: a review”. In: Ecological Entomology 46.2 (2021), pp. 143–159.

[24] Benjamin ER Rubin, et al. “Social behaviour in bees influences the abundance of Sodalis (Enterobacteriaceae) symbionts”. In: Royal Society Open Science 5.7 (2018), p. 180369.

[25] Wyatt A Shell and Sandra M Rehan. “Comparative metagenomics reveals expanded insights into intra-and interspecific variation among wild bee microbiomes”. In: Communications biology 5.1 (2022), p. 603.

[26] Quinn S McFrederick, et al. “Specificity between lactobacilli and hymenopteran hosts is the exception rather than the rule”. In: Applied and Environmental Microbiology 79.6 (2013), pp. 1803–1812.

[27] Sarah D Kocher and Robert J Paxton. “Comparative methods offer powerful insights into social evolution in bees”. In: Apidologie 45 (2014), pp. 289–305.

[28] Sean G Brady, et al. “Recent and simultaneous origins of eusociality in halictid bees”. In: Proceedings of the Royal Society B: Biological Sciences 273.1594 (2006), pp. 1643–1649.

[29] Jason Gibbs et al. “Phylogeny of halictine bees supports a shared origin of eusociality for Halictus and Lasioglossum (Apoidea: Anthophila: Halictidae)”. In: Molecular phylogenetics and evolution 65.3 (2012), pp. 926–939.

[30] Babraham Bioinformatics. “FastQC: a quality control tool for high throughput sequence data”. In: Cambridge, UK: Babraham Institute (2011).

[31] Philip Ewels, et al. “MultiQC: summarize analysis results for multiple tools and samples in a single report”. In: Bioinformatics 32.19 (2016), pp. 3047–3048.

[32] Anthony M Bolger, Marc Lohse, and Bjoern Usadel. “Trimmomatic: a flexible trimmer for Illumina sequence data”. In: Bioinformatics 30.15 (2014), pp. 2114–2120.

[33] Daehwan Kim, et al. “Graph-based genome alignment and genotyping with HISAT2 and HISAT-genotype”. In: Nature biotechnology 37.8 (2019), pp. 907–915.

[34] Beryl M Jones, et al. “Convergent and complementary selection shaped gains and losses of eusociality in sweat bees”. In: Nature Ecology & Evolution (2023), pp. 1–13.

[35] Olga Dudchenko, et al. “De novo assembly of the Aedes aegypti genome using Hi-C yields chromosome-length scaffolds”. In: Science 356.6333 (2017), pp. 92–95.

[36] Olga Dudchenko, et al. “The Juicebox Assembly Tools module facilitates de novo assembly of mammalian genomes with chromosome-length scaffolds for under $ @articledudchenko2018juicebox, title=The Juicebox Assembly Tools module facilitates de novo assembly of mammalian genomes with chromosome-length scaffolds for under $ 1000, author=Dudchenko, Olga and Shamim, Muhammad S and Batra, Sanjit S and Durand, Neva C and Musial, Nathaniel T and Mostofa, Ragib and Pham, Melanie and Glenn St Hilaire, Brian and Yao, Weijie and Stamenova, Elena and others, journal=BioRxiv, pages=254797, year=2018, publisher=Cold Spring Harbor Laboratory 1000”. In: BioRxiv (2018), p. 254797.

[37] Dmitry Meleshko, Iman Hajirasouliha, and Anton Korobeynikov. “coronaSPAdes: from biosynthetic gene clusters to RNA viral assemblies”. In: Bioinformatics 38.1 (2022), pp. 1–8.

[38] Stephen Nayfach, et al. “CheckV assesses the quality and completeness of metagenome-assembled viral genomes”. In: Nature biotechnology 39.5 (2021), pp. 578–585.

[39] Michael J Tisza, et al. “Cenote-Taker 2 democratizes virus discovery and sequence annotation”. In: Virus evolution 7.1 (2021), veaa100.

[40] Artem Babaian and Robert Edgar. “Ribovirus classification by a polymerase barcode sequence”. In: PeerJ 10 (2022), e14055.

[41] Limin Fu, et al. “CD-HIT: accelerated for clustering the next-generation sequencing data”. In: Bioinformatics 28.23 (2012), pp. 3150–3152.

[42] Christiam Camacho, et al. “BLAST+: architecture and applications”. In: BMC bioinformatics 10 (2009), pp. 1–9.

[43] Kim D Pruitt, Tatiana Tatusova, and Donna R Maglott. “NCBI reference sequences (RefSeq): a curated non-redundant sequence database of genomes, transcripts and proteins”. In: Nucleic acids research 35.suppl 1 (2007), pp. D61–D65.

[44] Eugene V Koonin, Valerian V Dolja, and T Jack Morris. “Evolution and taxonomy of positive-strand RNA viruses: implications of comparative analysis of amino acid sequences”. In: Critical reviews in biochemistry and molecular biology 28.5 (1993), pp. 375–430.

[45] Kazutaka Katoh and Daron M Standley. “MAFFT multiple sequence alignment software version 7: improvements in performance and usability”. In: Molecular biology and evolution 30.4 (2013), pp. 772–780.

[46] Salvador Capella-Gutiérrez, José M Silla-Martınez, and Toni Gabaldón. “trimAl: a tool for automated alignment trimming in large-scale phylogenetic analyses”. In: Bioinformatics 25.15 (2009), pp. 1972–1973.

[47] Bui Quang Minh, et al. “IQ-TREE 2: new models and efficient methods for phylogenetic inference in the genomic era”. In: Molecular biology and evolution 37.5 (2020), pp. 1530–1534.

[48] Subha Kalyaanamoorthy, et al. “ModelFinder: fast model selection for accurate phylogenetic estimates”. In: Nature methods 14.6 (2017), pp. 587–589.

[49] Heng Li. “Aligning sequence reads, clone sequences and assembly contigs with BWA-MEM”. In: arXiv preprint arXiv:1303.3997 (2013).

[50] Simon Roux, et al. “Benchmarking viromics: an in silico evaluation of metagenome-enabled estimates of viral community composition and diversity”. In: PeerJ 5 (2017), e3817.

[51] Ilias Lagkouvardos, et al. “Rhea: a transparent and modular R pipeline for microbial profiling based on 16S rRNA gene amplicons”. In: PeerJ 5 (2017), e2836.

[52] Felipe de Mendiburu and Maintainer Felipe de Mendiburu. “Package ‘agricolae’”. In: R Package, version 1.3 (2019).

[53] Jari Oksanen, et al. “The vegan package”. In: Community ecology package 10.631-637 (2007), p. 719.

[54] Hadley Wickham. “ggplot2”. In: Wiley interdisciplinary reviews: computational statistics 3.2 (2011), pp. 180–185.

[55] Andreas Wilm, et al. “LoFreq: a sequence-quality aware, ultra-sensitive variant caller for uncovering cell-population heterogeneity from high-throughput sequencing datasets”. In: Nucleic acids research 40.22 (2012), pp. 11189–11201.

[56] Ann C Gregory, et al. “Marine DNA viral macro-and microdiversity from pole to pole”. In: Cell 177.5 (2019), pp. 1109–1123.

[57] Eugene V Ryabov. “Invertebrate RNA virus diversity from a taxonomic point of view”. In: Journal of invertebrate pathology 147 (2017), pp. 37–50.

[58] Go Atsumi et al. “A novel virus transmitted through pollination causes ring-spot disease on gentian (Gentiana triflora) ovaries”. In: Journal of General Virology 96.2 (2015), pp. 431–439.

[59] Michael J Adams et al. “ICTV virus taxonomy profile: Virgaviridae”. In: The Journal of general virology 98.8 (2017), p. 1999.

[60] Humberto J Debat. “An RNA virome associated to the golden orb-weaver spider Nephila clavipes”. In: Frontiers in microbiology 8 (2017), p. 2097.

[61] Minh Vong, et al. “Complete cryspovirus genome sequences from Cryptosporidium parvum isolate Iowa”. In: Archives of virology 162 (2017), pp. 2875–2879.

[62] SM Valles et al. “ICTV virus taxonomy profile: Iflaviridae”. In: The Journal of general virology 98.4 (2017), p. 527.

[63] Stephen J Martin and Laura E Brettell. “Deformed wing virus in honeybees and other insects”. In: Annual review of virology 6 (2019), pp. 49–69.

[64] Lena Wilfert, et al. “Deformed wing virus is a recent global epidemic in honeybees driven by Varroa mites”. In: Science 351.6273 (2016), pp. 594–597.

[65] Laura E Brettell, et al. “A comparison of deformed wing virus in deformed and asymptomatic honey bees”. In: Insects 8.1 (2017), p. 28.

[66] Robyn Manley, Mike Boots, and Lena Wilfert. “Condition-dependent virulence of slow bee paralysis virus in Bombus terrestris: are the impacts of honeybee viruses in wild pollinators underestimated?” In: Oecologia 184 (2017), pp. 305–315.

[67] Kishor Dhaygude, et al. “Genome organization and molecular characterization of the three Formica exsecta viruses—FeV1, FeV2 and FeV4”. In: PeerJ 6 (2019), e6216.

[68] Nolwenn M Dheilly, et al. “Who is the puppet master? Replication of a parasitic wasp-associated virus correlates with host behaviour manipulation”. In: Proceedings of the Royal Society B: Biological Sciences 282.1803 (2015), p. 20142773.

[69] Bradley I Hillman and Guohong Cai. “The family Narnaviridae: simplest of RNA viruses”. In: Advances in virus research 86 (2013), pp. 149–176.

[70] Masatoshi Nei. “Analysis of gene diversity in subdivided populations”. In: Proceedings of the national academy of sciences 70.12 (1973), pp. 3321–3323.

[71] Vincent G Martinson, et al. “A simple and distinctive microbiota associated with honey bees and bumble bees”. In: Molecular ecology 20.3 (2011), pp. 619–628.

[72] Karen M Kapheim, Makenna M Johnson, and Maggi Jolley. “Composition and acquisition of the microbiome in solitary, ground-nesting alkali bees”. In: Scientific reports 11.1 (2021), p. 2993.

[73] Waldan K Kwong and Nancy A Moran. “Gut microbial communities of social bees”. In: Nature reviews microbiology 14.6 (2016), pp. 374–384.

[74] Jo-Anne C Holley, et al. “Carpenter Bees (Xylocopa) harbor a distinctive gut microbiome related to that of honey bees and bumble bees”. In: Applied and Environmental Microbiology 88.13 (2022), e00203–22.

[75] Charles Darwin. On the origin of species, 1859. 2016.

