## Supplementary material for "Host species and geography impact bee-associated RNA virus communities with evidence for isolation-by-distance in viral populations": Suupelemental Data

### 1 Supplemental Figures

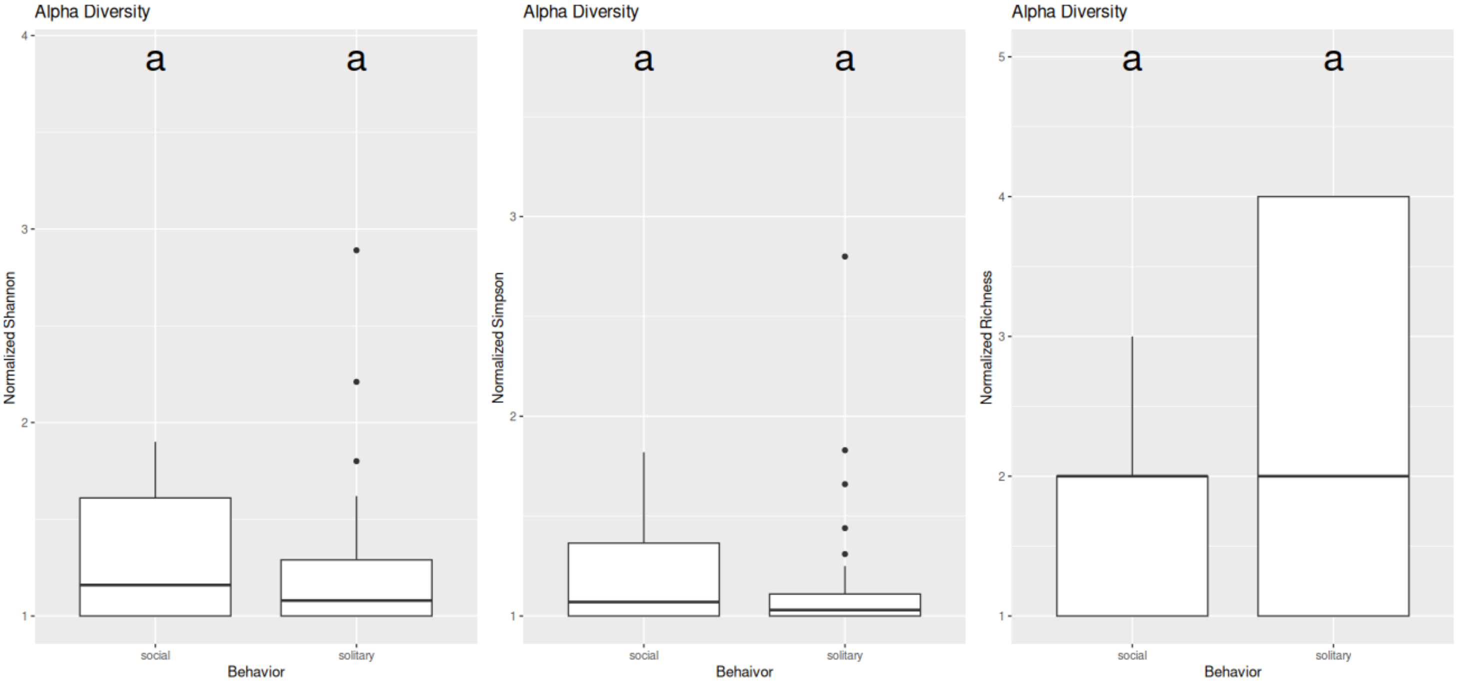


Figure 1: Alpha diversity metrics (Shannon index, Simpson index, and Richness) describing differences in viral species diversity associated with social and solitary bees.

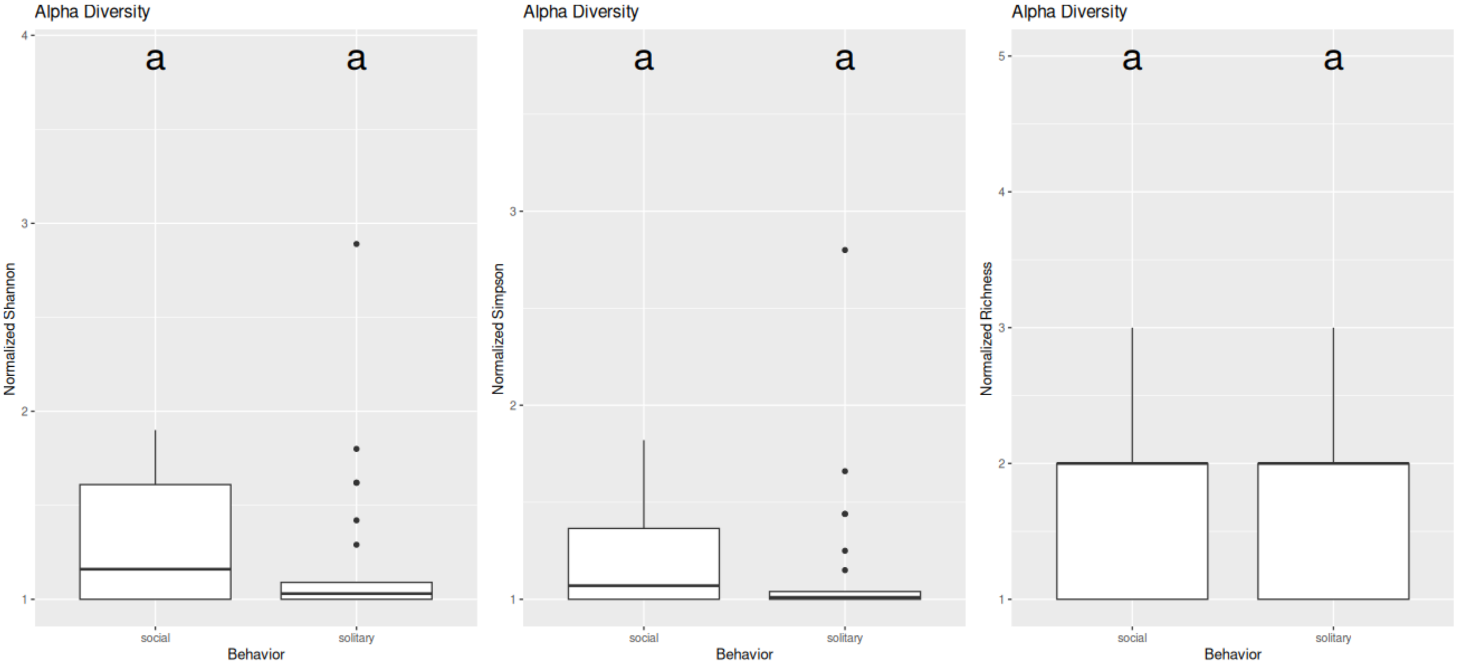


Figure 2: Alpha diversity metrics (Shannon index, Simpson index, and Richness) describing differences in viral family diversity associated with social and solitary bees. Letters correspond to groupings by Tukey's Honest Significant Difference Test

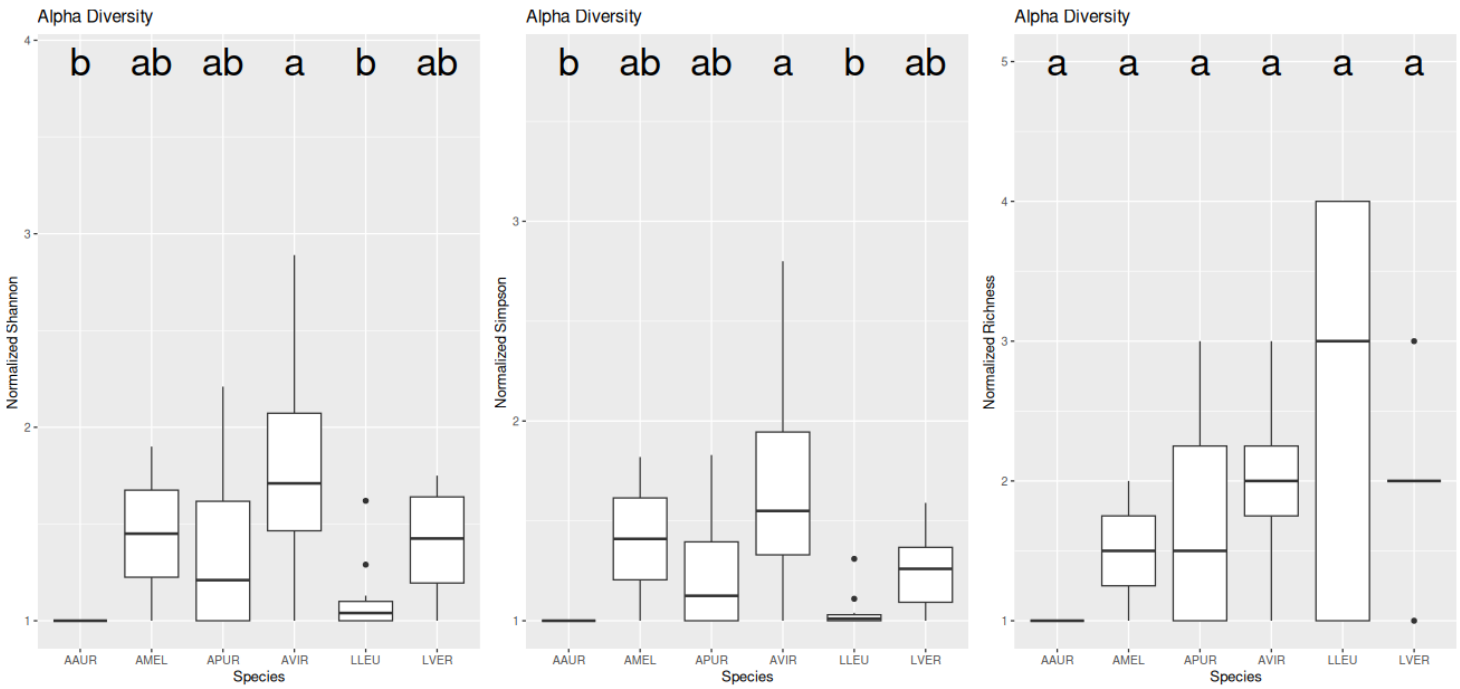


Figure 3: Alpha diversity metrics (Shannon index, Simpson index, and Richness) describing differences in viral species diversity associated with different bee species. Letters correspond to groupings by Tukey's Honest Significant Difference Test

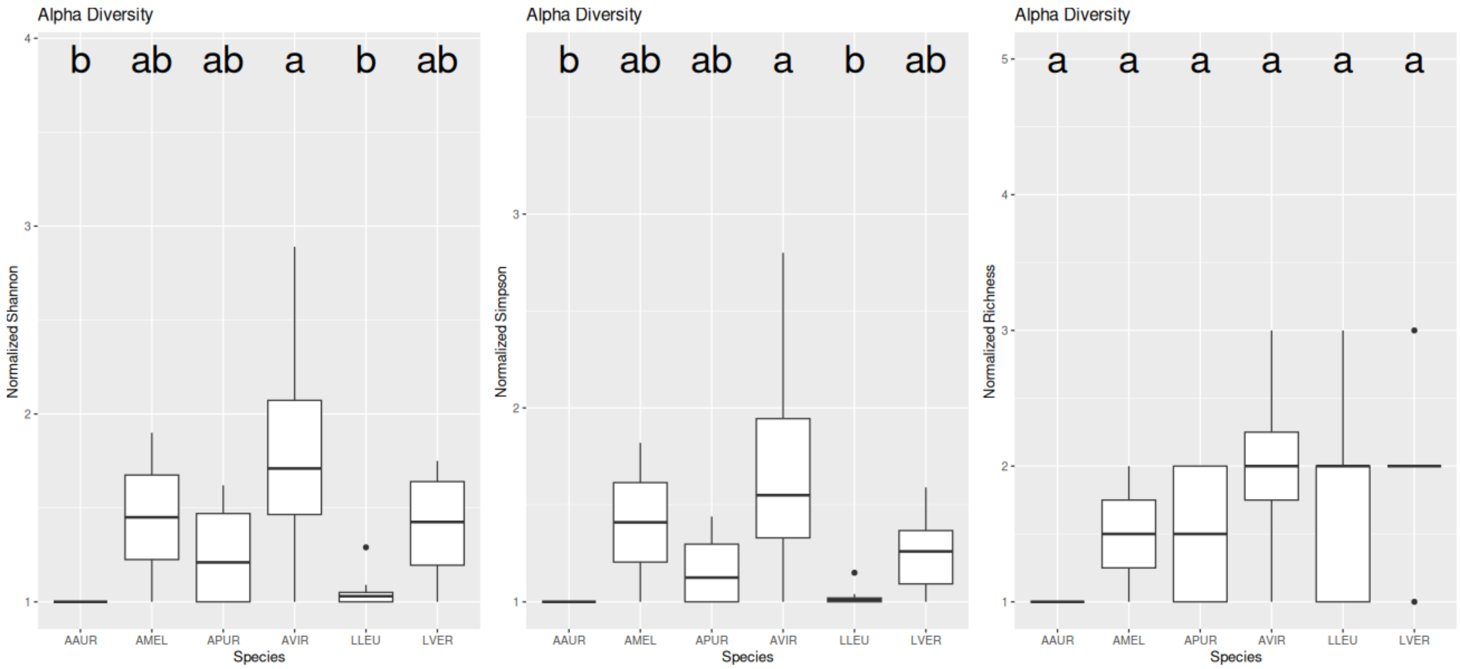


Figure 4: Alpha diversity metrics (Shannon index, Simpson index, and Richness) describing differences in viral family diversity associated with different bee species. Letters correspond to groupings by Tukey's Honest Significant Difference Test.

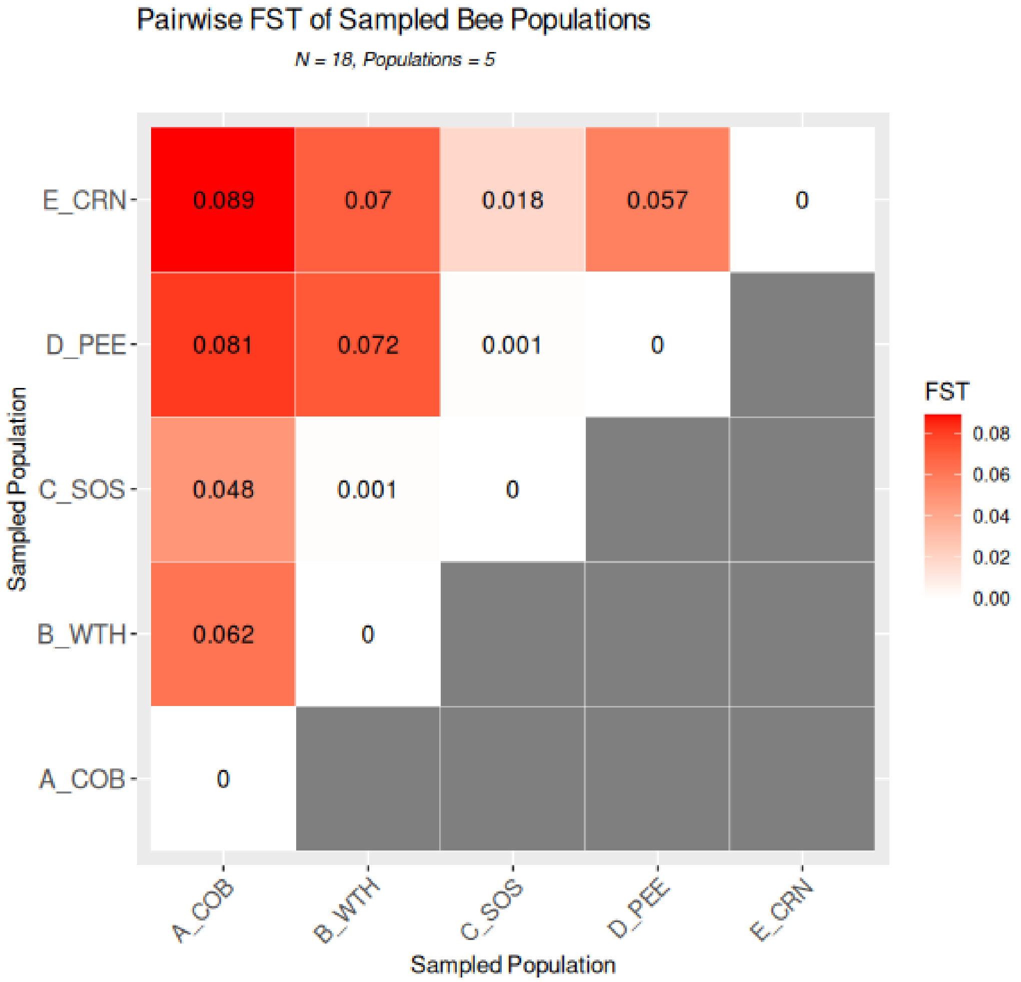


Figure 5: A matrix of pairwise $F_{st}$ comparisons representing patterns of genetic differentiation among Narnaviridae populations associated with 5 geographically-defined sites. Color represents the degree of genetic differentiation between any pair of combinations. Darker color indicates higher values of differentiation. Genetic differentiation is defined by a scale on the right side of the figure.

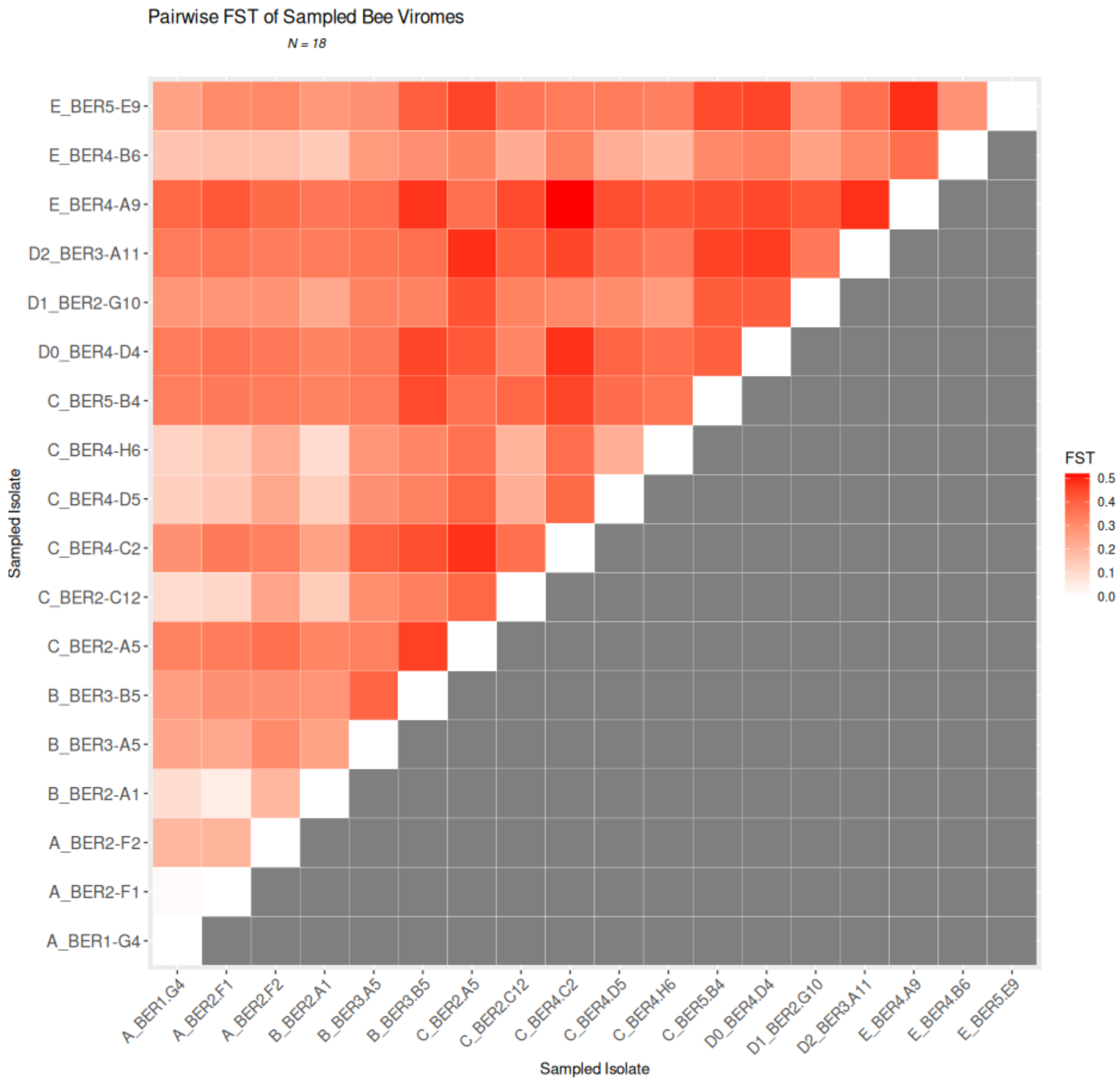


Figure 6: A matrix of pairwise $F_{st}$ comparisons representing patterns of genetic differentiation among 18 individual populations of *Narnaviridae* viromes. Color represents the degree of genetic differentiation between any pair of combinations. Darker color indicates higher values of differentiation. Genetic differentiation is defined by a scale on the right side of the figure.

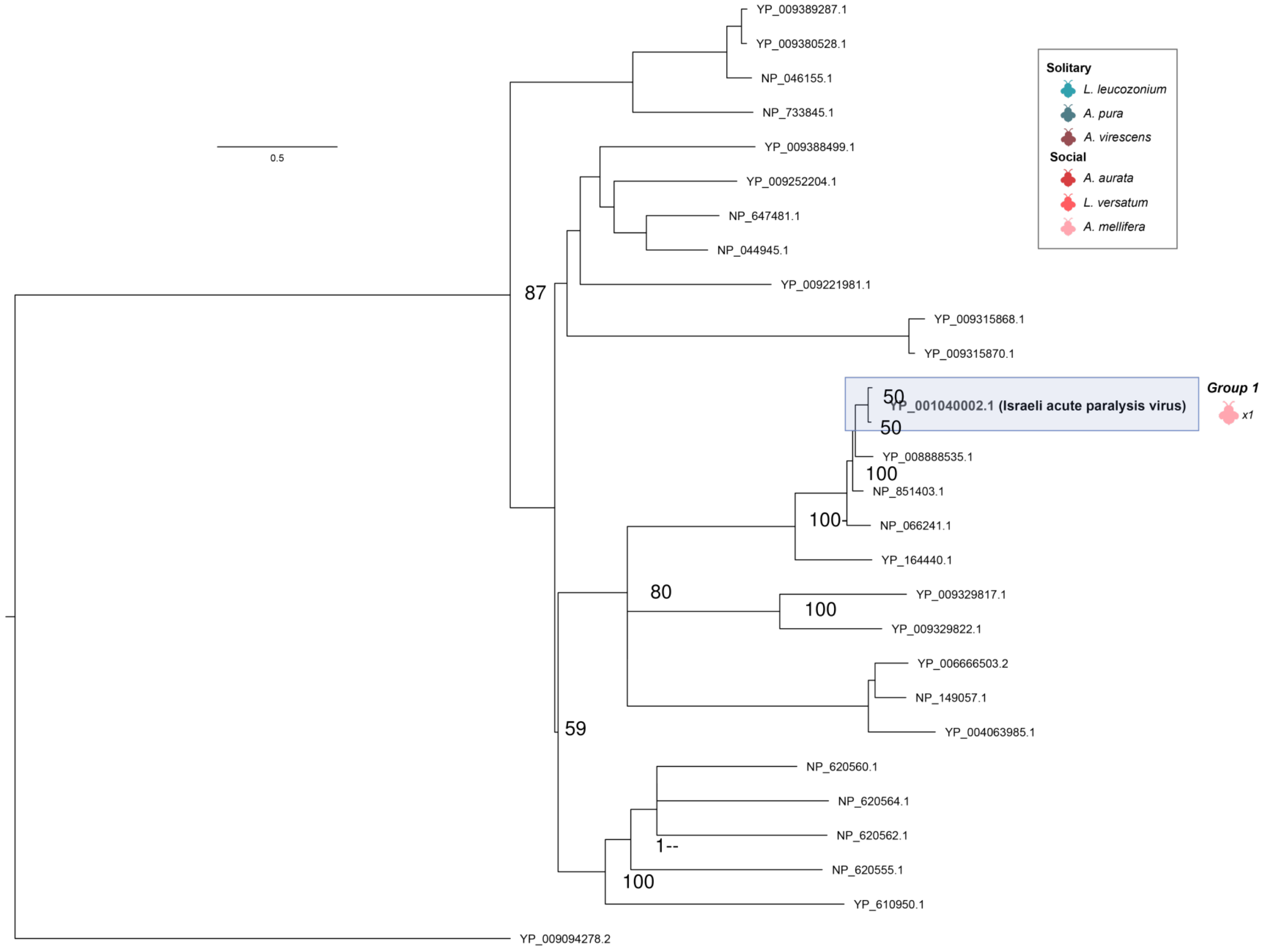


Figure 7: RdRP phylogeny of Dicistroviridae viruses. Tree is midpoint rooted for clarity only. Viruses sampled in this study are detailed highlighted in blue. Bootstrap values for select nodes are provided. Bee icon color represent species host that viral group was associated with; number represent the number of bee samples associated with viral group. The scale bar represents the number of amino acid substitutions.

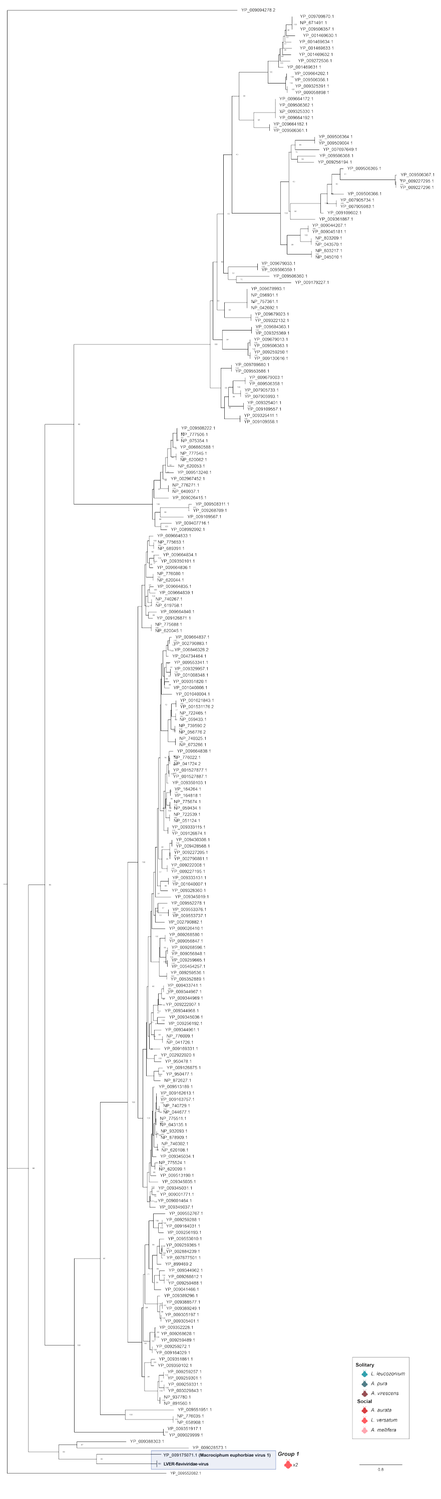


Figure 8: RdRP phylogeny of Flaviviridae viruses. Tree is midpoint rooted for clarity only. Viruses sampled in this study are detailed highlighted in blue. Bootstrap values for select nodes are provided. Bee icon color represent species host that viral group was associated with; number represent the number of bee samples associated with viral group. The scale bar represents the number of amino acid substitutions.

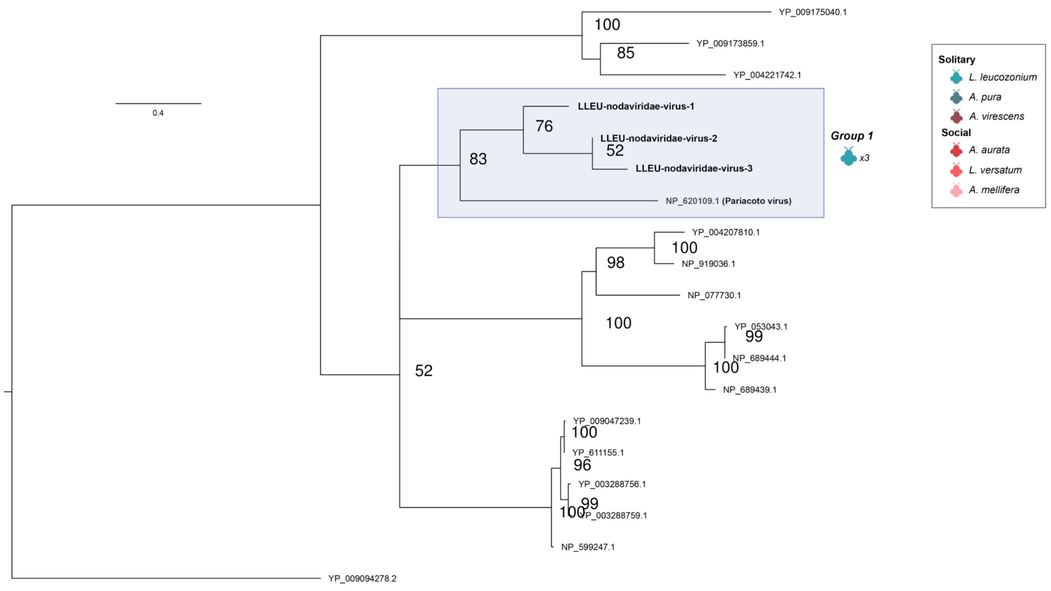


Figure 9: RdRP phylogeny of Nodaviridae viruses. Tree is midpoint rooted for clarity only. Viruses sampled in this study are detailed highlighted in blue. Bootstrap values for select nodes are provided. Bee icon color represent species host that viral group was associated with; number represent the number of bee samples associated with viral group. The scale bar represents the number of amino acid substitutions.

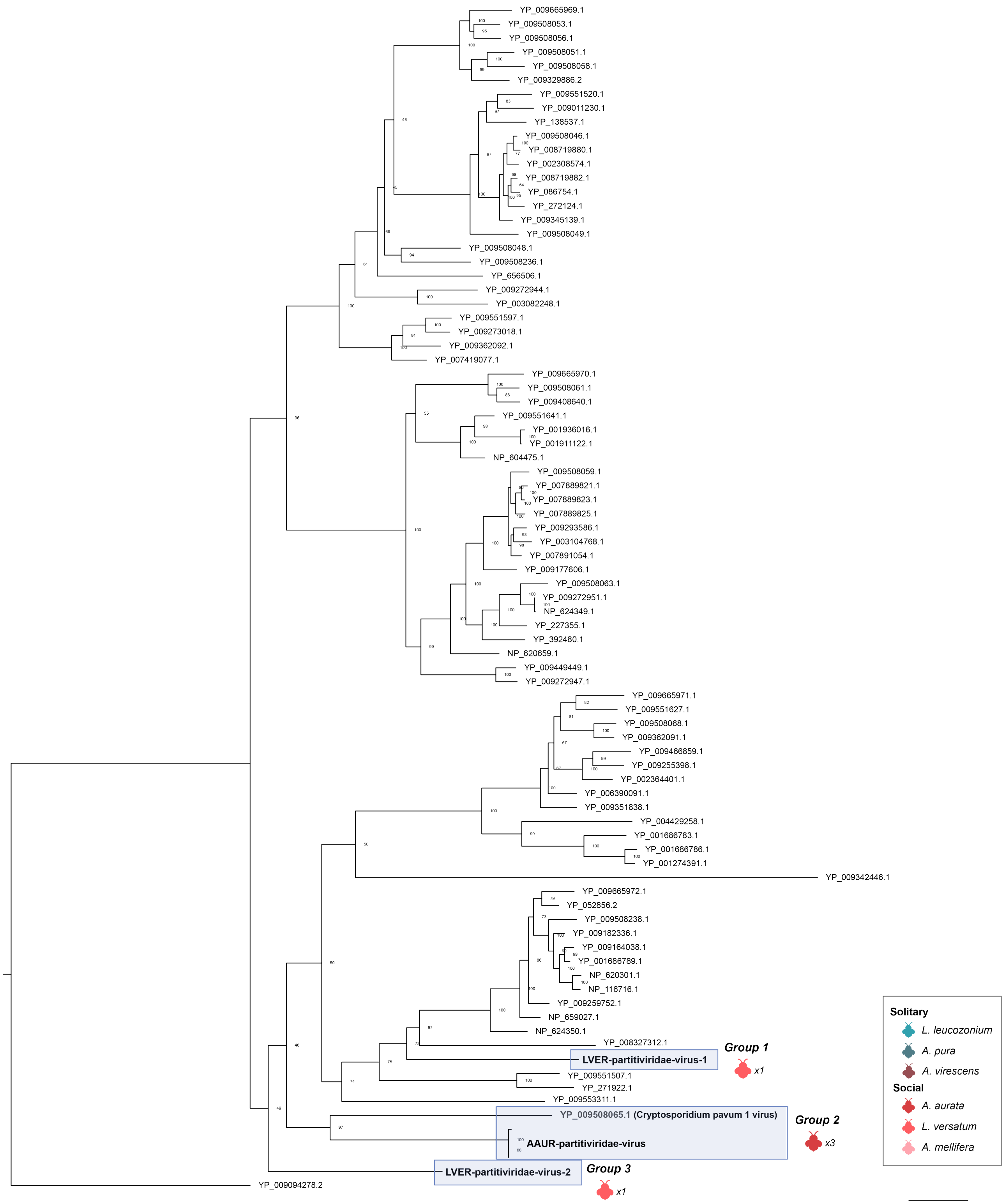


Figure 10: RdRP phylogeny of Partitiviridae viruses. Tree is midpoint rooted for clarity only. Viruses sampled in this study are detailed highlighted in blue. Bootstrap values for select nodes are provided. Bee icon color represent species host that viral group was associated with; number represent the number of bee samples associated with viral group. The scale bar represents the number of amino acid substitutions.

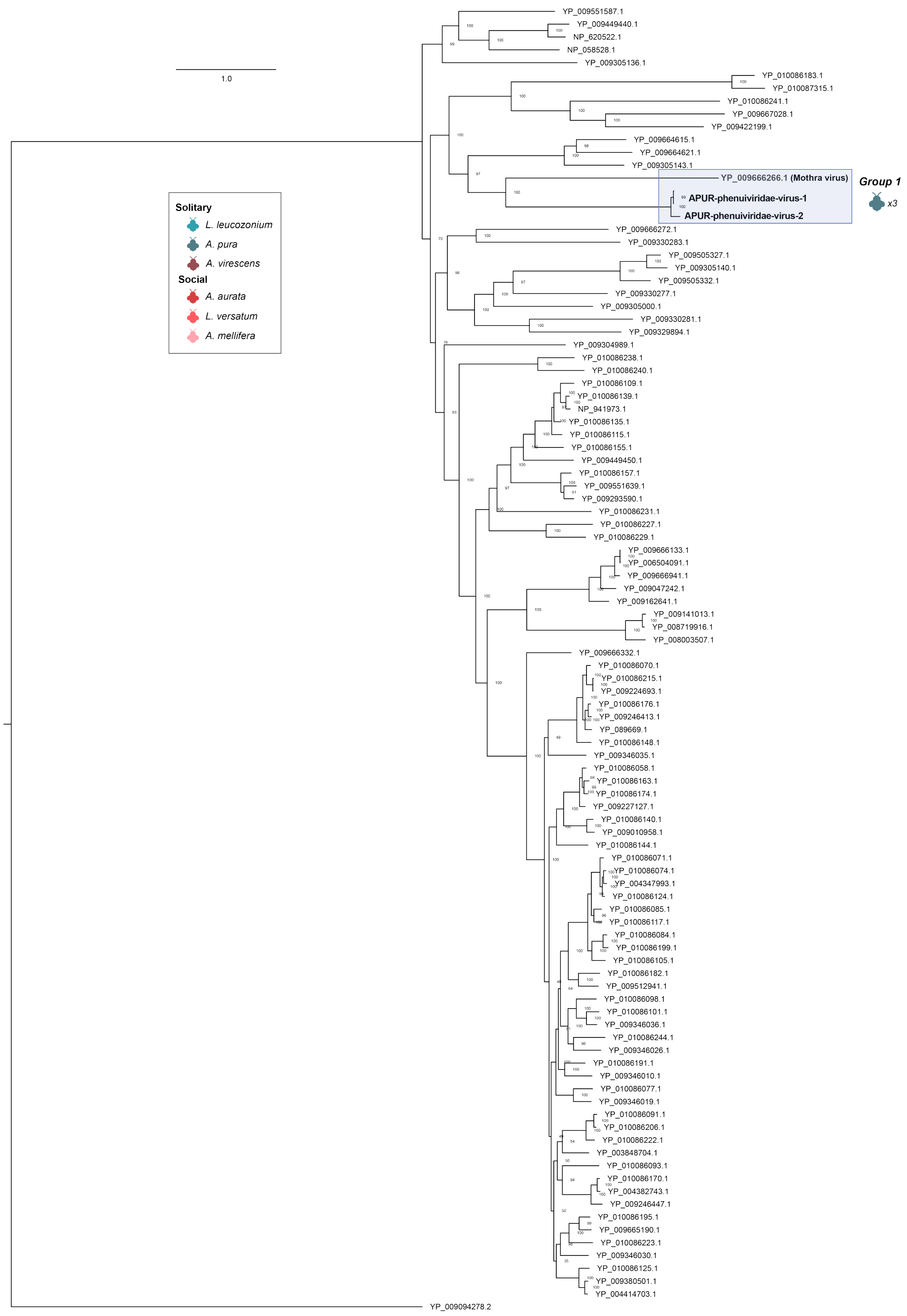


Figure 11: RdRP phylogeny of Phenuiviridae viruses. Tree is midpoint rooted for clarity only. Viruses sampled in this study are detailed highlighted in blue. Bootstrap values for select nodes are provided. Bee icon color represent species host that viral group was associated with; number represent the number of bee samples associated with viral group. The scale bar represents the number of amino acid substitutions.

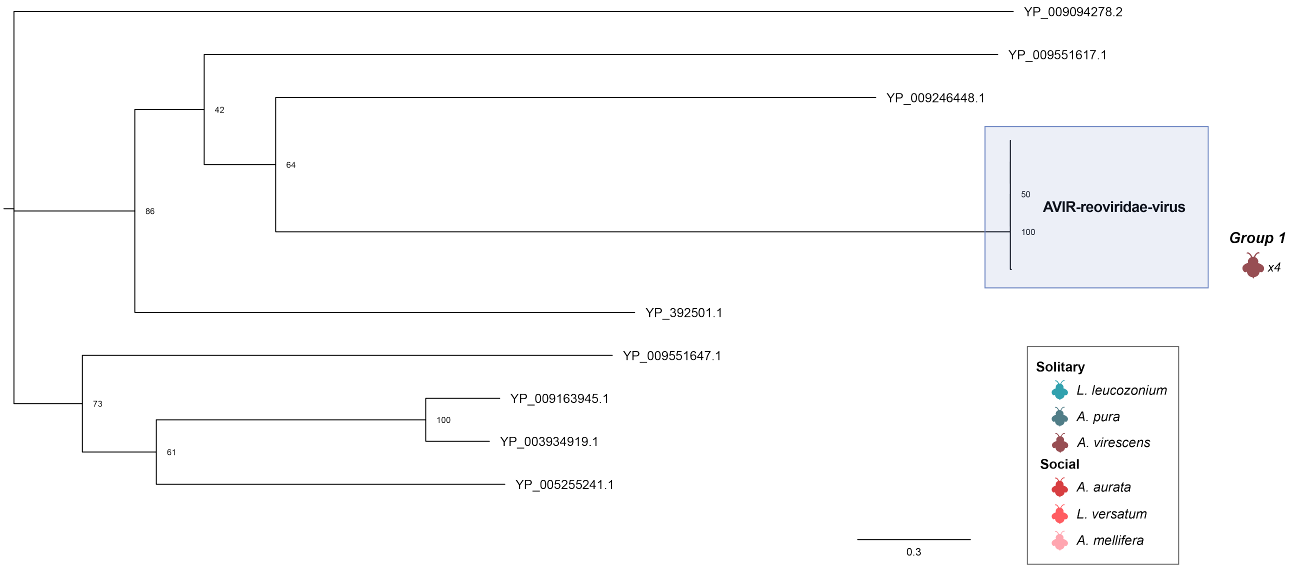


Figure 12: RdRP phylogeny of Reoviridae viruses. Tree is midpoint rooted for clarity only. Viruses sampled in this study are detailed highlighted in blue. Bootstrap values for select nodes are provided. Bee icon color represent species host that viral group was associated with; number represent the number of bee samples associated with viral group. The scale bar represents the number of amino acid substitutions.

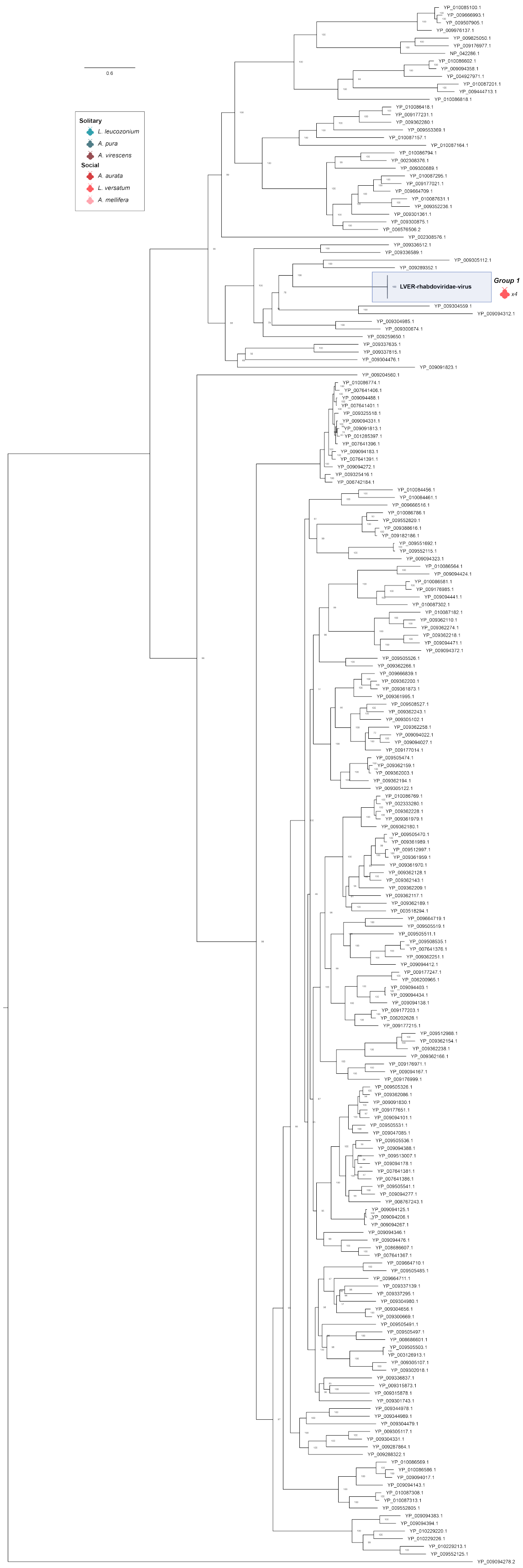


Figure 13: RdRP phylogeny of Rhabodiviridae viruses. Tree is midpoint rooted for clarity only. Viruses sampled in this study are detailed highlighted in blue. Bootstrap values for select nodes are provided. Bee icon color represent species host that viral group was associated with; number represent the number of bee samples associated with viral group. The scale bar represents the number of amino acid substitutions.

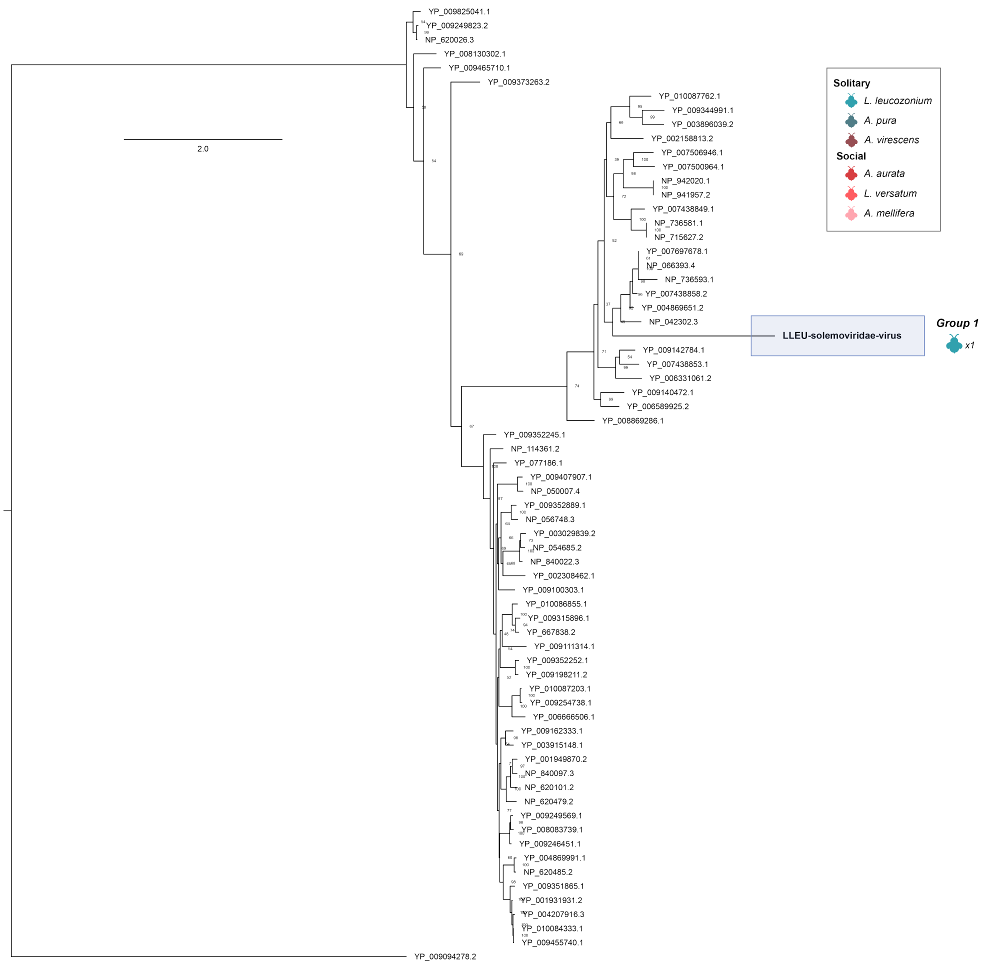


Figure 14: RdRP phylogeny of Solemoviridae viruses. Tree is midpoint rooted for clarity only. Viruses sampled in this study are detailed highlighted in blue. Bootstrap values for select nodes are provided. Bee icon color represent species host that viral group was associated with; number represent the number of bee samples associated with viral group. The scale bar represents the number of amino acid substitutions.

| Sample | Sex | Site | Behavior | Latitude | Longitude | Genus | Species | DateCollected |
|---|---|---|---|---|---|---|---|---|
| BER1-G4 | M | Cobscook Bay, ME | solitary | 44.83916 | 67.1504 | Lasioglossum | leucozonium | Jul2016 |
| BER2-A1 | F | Winter Harbor, ME | solitary | 44.39461 | 68.0846 | Lasioglossum | leucozonium | Jul2016 |
| BER2-A5 | F | Rangley, ME | solitary | 44.92854 | 70.6362 | Lasioglossum | leucozonium | Jul2016 |
| BER2-C12 | F | Rangley, ME | solitary | 44.92854 | 70.6362 | Lasioglossum | leucozonium | Jul2016 |
| BER2-F1 | M | Cobscook Bay, ME | solitary | 44.83916 | 67.1504 | Lasioglossum | leucozonium | Jul2016 |
| BER2-F2 | M | Cobscook Bay, ME | solitary | 44.83916 | 67.1504 | Lasioglossum | leucozonium | Jul2016 |
| BER2-G10 | M | Sunapee, NH | solitary | 43.38236 | 72.0854 | Lasioglossum | leucozonium | Jul2016 |
| BER3-A11 | M | Sunapee, NH | solitary | 43.38236 | 72.0854 | Lasioglossum | leucozonium | Jul2016 |
| BER3-A5 | F | Winter Harbor, ME | solitary | 44.39461 | 68.0846 | Lasioglossum | leucozonium | Jul2016 |
| BER3-B5 | F | Winter Harbor, ME | solitary | 44.39461 | 68.0846 | Lasioglossum | leucozonium | Jul2016 |
| BER4-A9 | F | Cranberry Lake, NY | solitary | 44.20406 | 74.8312 | Lasioglossum | leucozonium | Jul2016 |
| BER4-B6 | F | Cranberry Lake, NY | solitary | 44.20406 | 74.8312 | Lasioglossum | leucozonium | Jul2016 |
| BER4-C2 | F | Rangley, ME | solitary | 44.92854 | 70.6362 | Lasioglossum | leucozonium | Jul2016 |
| BER4-D4 | M | Sunapee, NH | solitary | 43.38236 | 72.0854 | Lasioglossum | leucozonium | Jul2016 |
| BER4-D5 | M | Rangley, ME | solitary | 44.92854 | 70.6362 | Lasioglossum | leucozonium | Jul2016 |
| BER4-H6 | M | Rangley, ME | solitary | 44.92854 | 70.6362 | Lasioglossum | leucozonium | Jul2016 |
| BER5-B4 | M | Rangley, ME | solitary | 44.92854 | 70.6362 | Lasioglossum | leucozonium | Jul2016 |
| BER5-E9 | F | Cranberry Lake, NY | solitary | 44.20406 | 74.8312 | Lasioglossum | leucozonium | Jul2016 |
| BMJ2-B12 | F | Princeton, NJ | solitary | 40.33707 | 74.6526 | Augochlora | pura | Sep2018 |
| BMJ2-B6 | M | Princeton, NJ | solitary | 40.33707 | 74.6526 | Augochlora | pura | Sep2018 |
| BMJ2-D10 | M | Princeton, NJ | social | 40.33707 | 74.6526 | Augochlora | pura | Sep2018 |
| BMJ2-D11 | F | Princeton, NJ | solitary | 40.33707 | 74.6526 | Augochlora | pura | Sep2018 |
| BMJ2-D2 | M | Princeton, NJ | social | 40.33707 | 74.6526 | Lasioglossum | versatum | Sep2018 |
| BMJ2-E12 | F | Princeton, NJ | social | 40.33707 | 74.6526 | Augochlorella | aurata | Sep2018 |
| BMJ2-E2 | F | Princeton, NJ | social | 40.33707 | 74.6526 | Augochlorella | aurata | Sep2018 |
| BMJ2-F6 | F | Princeton, NJ | solitary | 40.33707 | 74.6526 | Agapostemon | virescens | Sep2018 |
| BMJ2-F7 | F | Princeton, NJ | solitary | 40.33707 | 74.6526 | Agapostemon | virescens | Sep2018 |
| BMJ2-G12 | F | Princeton, NJ | social | 40.33707 | 74.6526 | Augochlorella | aurata | Sep2018 |
| BMJ3-A9 | M | Princeton, NJ | social | 40.33707 | 74.6526 | Lasioglossum | versatum | Sep2018 |
| BMJ3-B1 | F | Princeton, NJ | social | 40.33707 | 74.6526 | Lasioglossum | versatum | Sep2018 |
| BMJ3-B10 | F | Princeton, NJ | social | 40.33707 | 74.6526 | Lasioglossum | versatum | Sep2018 |
| BMJ3-B11 | M | Princeton, NJ | social | 40.33707 | 74.6526 | Lasioglossum | versatum | Sep2018 |
| BMJ3-B3 | F | Princeton, NJ | social | 40.33707 | 74.6526 | Lasioglossum | versatum | Sep2018 |
| BMJ3-F5 | F | Princeton, NJ | solitary | 40.33707 | 74.6526 | Agapostemon | virescens | Sep2018 |
| BMJ3-F9 | F | Princeton, NJ | honey | 40.33707 | 74.6526 | Apis | mellifera | Sep2018 |
| BMJ3-H11 | M | Princeton, NJ | solitary | 40.33707 | 74.6526 | Agapostemon | virescens | Sep2018 |
| BMJ3-H3 | F | Princeton, NJ | honey | 40.33707 | 74.6526 | Apis | mellifera | Sep2018 |
| BMJ3-H9 | F | Princeton, NJ | solitary | 40.33707 | 74.6526 | Augochlora | aurata | Sep2018 |

Table 1: Metadata associated with sampled bees.
